# Dynamical phase diagram of an auto-regulating gene in fast switching conditions

**DOI:** 10.1101/2020.03.10.985291

**Authors:** Chen Jia, Ramon Grima

## Abstract

While the steady-state behavior of stochastic gene expression with auto-regulation has been extensively studied, its time-dependent behavior has received much less attention. Here, under the assumption of fast promoter switching, we derive and solve a reduced chemical master equation for an auto-regulatory gene circuit with translational bursting and cooperative protein-gene interactions. The analytical expression for the time-dependent probability distribution of protein numbers enables a fast exploration of large swaths of parameter space. For a unimodal initial distribution, we identify three distinct types of stochastic dynamics: (i) the protein distribution remains unimodal at all times; (ii) the protein distribution becomes bimodal at intermediate times and then reverts back to being unimodal at long times (transient bimodality) and (iii) the protein distribution switches to being bimodal at long times. For each of these, the deterministic model predicts either monostable or bistable behaviour and hence there exist six dynamical phases in total. We investigate the relationship of the six phases to the transcription rates, the protein binding and unbinding rates, the mean protein burst size, the degree of cooperativity, the relaxation time to the steady state, the protein mean and the type of feedback loop (positive or negative). We show that transient bimodality is a noise-induced phenomenon that occurs when protein expression is sufficiently bursty and we use theory to estimate the observation time window when it is manifest.

## 1 Introduction

Auto-regulation, whereby the protein expressed from a gene activates or suppresses its own transcription, is the most common form of feedback in gene regulatory systems [1]. Experimental studies have probed how feedback loops modulate fluctuations in the concentrations of gene products [2–4]. The exact distribution of protein numbers in four distinct stochastic models of auto-regulation [5–8] has been obtained by solution of the chemical master equation (CME, [9]) in steady-state conditions. These four models do not consider cooperativity of protein binding to the gene and are based on different implicit assumptions, as follows. Hornos et al. [5] and Kumar et al. [6] neglect binding fluctuations while Grima et al. [7] and Jia et al. [8] take them into account. Note that by binding fluctuations here we mean a decrease (increase) in protein number whenever a binding (unbinding) reaction occurs. In addition, Hornos et al. and Grima et al. assume non-bursty protein expression while Kumar et. al and Jia et al. take into account bursty expression. Since protein expression is often bursty and since binding fluctuations can be important for cases where the protein is in low numbers, it follows that the model in [8] is the most realistic among the four. In all cases, the steady-state protein distributions can be either unimodal or bimodal even though the corresponding deterministic model is always monostable (since there is no cooperativity). Slow switching between the ON and OFF states of the gene can lead to bimodality if the transcription rates in the two gene states are well separated, for both positive and negative feedback. Fast switching often leads to unimodality but under certain conditions, a positive feedback loop can also produce bimodality with one of the modes approximately centered on zero [8]. For a recent review of these and other similar models, see [10]. For recently developed methods to approximately solve for the steady-state distribution in models of auto-regulation, see [11–13].

The time-dependent solution for stochastically modelled chemical reaction systems has received comparatively very little attention. Under certain conditions, for an initial joint distribution given by a product of Poissons, the transient joint distribution remains a product of Poissons for all times [9, 14]. However, the conditions for this result to hold are very restrictive and not applicable to most systems of biological relevance. To our knowledge, the only exact time-dependent solution for an auto-regulatory feedback loop is the one derived by Ramos et al. [15], where the authors obtain a Heun function for the generating function of a stochastic model neglecting translational bursting, cooperativity and binding fluctuations, i.e. the model of Hornos et al. [5]. An approximate time-dependent solution for four different types of auto-regulatory feedback loops was obtained by Cao and Grima using the linear mapping approximation [12], all of which have generating functions in terms of hypergeometric functions. Approximate time-dependent solutions have also been recently reported for detailed models of eukaryotic gene expressions with and without feedback and including binomial partitioning due to cell division [16]. Another method by Veerman et al. calculates the approximate time-dependent solution of auto-regulatory feedback loops by means of a perturbative approach [17]. However, the difficulty in analyzing Heun and hypergeometric functions means that little information can be extracted about the time-dependent behavior and hence different dynamical phases of auto-regulatory circuits still remain unknown. Elucidating and understanding such behavior is important since living cells are constantly exposed to an ever-changing environment that requires dynamic fine tuning of gene expression to maintain healthy cellular functions.

In this article, by means of careful approximations, we derive an analytical time-dependent solution for the CME of an auto-regulatory circuit in terms of a sum of simple functions and subsequently use it to study the phase diagram characterizing the dynamics of the gene circuit. The paper is divided as follows. In Section 2, starting from the CME of an auto-regulatory circuit with bursty protein expression in two gene states, cooperative protein-gene interactions and taking into account binding fluctuations, we use the method of multiscale averaging to obtain a reduced master equation valid in the limit of fast promoter switching. In Section 3, this reduced master equation is solved in time, leading to a time-dependent protein distribution that has the form of a sum of exponential functions; the analytical solution is found to be in excellent agreement with the numerical solution obtained using the finite state projection algorithm (FSP). In Section 4, we use the analytical solution to show that there are six different ways in which the time-dependent dynamics can unfold. The relationship of these dynamical phases to the values of model parameters, the relaxation time to the steady state and the mean protein number is investigated. The most interesting of these phases is transient bimodality, whereby the protein distribution is unimodal for short and long times but is bimodal for intermediate times; the theory is used to estimate the time window when such behavior can be observed. We conclude in Section 5.

## 2 Deriving a reduced model of auto-regulated bursty gene expression

We start by considering a stochastic model of auto-regulation that includes promoter switching, bursty protein expression, protein decay and feedback mediated by cooperative protein binding to the gene (see Fig. 1(a) for an illustration). Let *G* and *G*^∗^ denote the unbound and bound states of the gene, respectively, and let *P* denote the corresponding protein. The effective reactions describing the model are given by:

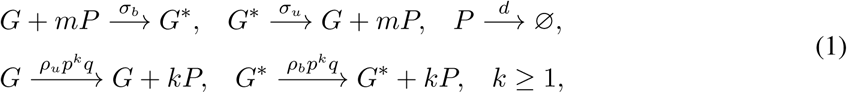

where *σ*_*b*_ is the binding rate of the protein to the promoter, *σ*_*u*_ is the unbinding rate of the protein from the promoter, *ρ*_*b*_ and *ρ*_*u*_ are the transcription rates when the protein is bound to or unbound from the promoter, respectively, and *d* is the decay rate of the protein due to active protein degradation and dilution during cell division [18]. The ratio *L* = *σ*_*b*_*/σ*_*u*_ of the protein binding and unbinding rates characterizes the strength of auto-regulation. In agreement with experiments [19, 20], protein production is assumed to occur in bursts of random size sampled from a geometric distribution with parameter *p*. Each burst is due to rapid translation of protein from a single, short-lived mRNA molecule; hence the mRNA dynamics is modelled implicitly. The effective translation rate in the unbound or bound gene state is then the product of the corresponding transcription rate, *ρ*_*u*_ or *ρ*_*b*_, and the geometric distribution *p*^*k*^*q*, where *q* = 1 − *p*. Since the protein burst size is geometrically distributed, its expected value is given by 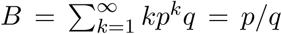 The reaction scheme describes a positive feedback loop if *ρ*_*b*_ *> ρ*_*u*_ and a negative feedback loop if *ρ*_*b*_ *< ρ*_*u*_. This model has been derived from a model that explicitly takes into account mRNA dynamics in [8]. Note that our model takes into account protein-gene binding fluctuations and hence as shown in [8] it is generally more realistic than the classical model of Kumar et al. [6], resulting in more accurate protein distributions for the case of low protein numbers.

**Fig. 1.**
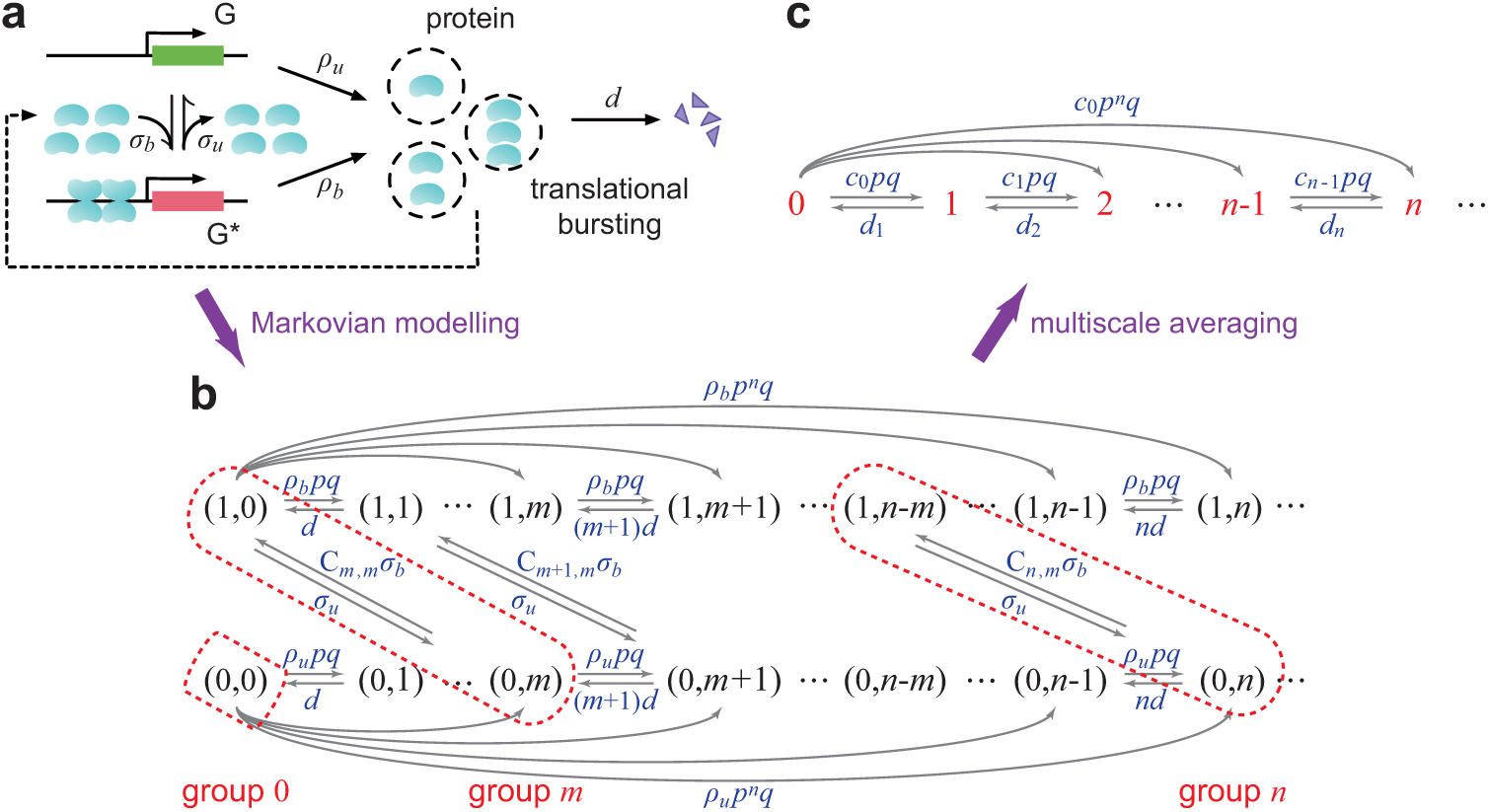
Autoregulated bursty gene expression with cooperative binding. (a) Schematic diagram of the stochastic model of auto-regulation. Here we depict the case when four protein molecules bind to the promoter cooperatively to regulate gene expression. (b) Markovian dynamics of the stochastic model of auto-regulation. When the gene switches rapidly between the unbound and bound states, for each *n* ≥ *m*, the two microstates (0, *n*) and (1, *n* − *m*) can be combined into a group that is labelled by group *n*. (c) Transition diagram of the reduced model in the limit of fast gene switching. When *n* ≥ *m*, group *n* is composed of two microstates with different protein numbers and thus the group index *n* cannot be interpreted as the protein number. Note that translational bursting can cause jumps from any group to another (this is shown for group 0 in the figure but is also true for any other groups).

The microstate of the gene of interest can be represented by an ordered pair (*i, n*): the state *i* of the gene with *i* = 0, 1 corresponding to the unbound and bound states, respectively, and the number *n* of the protein. Let *p*_*i,n*_ denote the probability of having *n* protein copies when the gene is in state *i*. Then the stochastic gene expression kinetics in a single cell can be described by the Markov jump process illustrated in Fig. 1(b). The evolution of the Markovian model is governed by the CME:

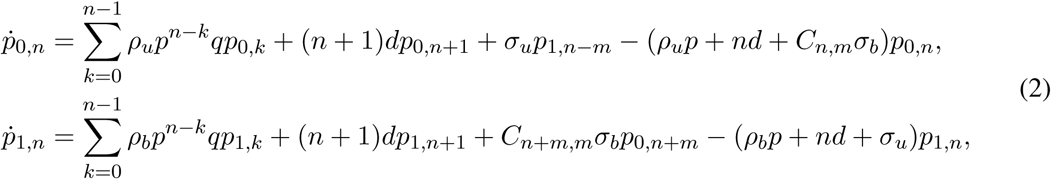

where *C*_*n,m*_ = *n*!*/m*!(*n* − *m*)! is the number of ways of choosing an unordered subset of *m* molecules from a set of *n* molecules. On the right-hand side of the first equation of the CME, the sum in the first term is taken from 0 to *n* − 1 because the burst size is ≥ 1. Note also that 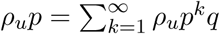 in the last term is the sum of all transition rates leaving microstate (0, *n*) due to random bursts of size ≥ 1. The corresponding terms in the second equation of the CME can be understood in the same way.

We next focus on the regime of fast gene switching, i.e. *σ*_*b*_, *σ*_*u*_ ≫ *ρ*_*b*_, *ρ*_*u*_, *d*. While this is a common simplifying assumption in many theoretical studies [21, 22], it is also supported by recent single-cell data in bacteria [23]. In this case, the Markovian model illustrated in Fig. 1(b) can be reduced to a much simpler one by using a classical simplification method of multiscale Markov jump processes called averaging [24–26]. *Since σ*_*b*_ *and σ*_*u*_ *are large, for each n* ≥ *m, the two microstates* (0, *n*) *and* (1, *n* − *m*) *are in rapid equilibrium and thus can be aggregated into a group that is labelled by group n*, as depicted in Fig. 1(b). In addition, for each *n < m*, group *n* is composed of the single microstate (0, *n*). In this way, the full Markovian model can be simplified to the reduced one shown in Fig. 1(c), whose state space is given by

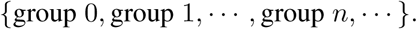

*Here we emphasize that the group index n cannot be interpreted as the protein number*. This is because when *n* ≥ *m*, group *n* is composed of two microstates with different protein numbers.

The next step is to determine the transition diagram and calculate the effective transition rates of the reduced model. In the fast switching limit, the two microstates (0, *n*) and (1, *n* − *m*) will reach a quasi-steady state with quasi-steady-state distribution

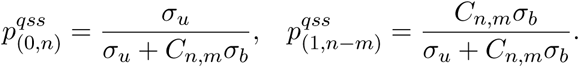

Then the effective transcription rate is given by

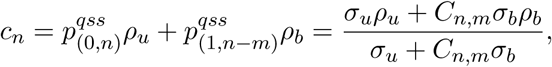

and the effective protein decay rate is given by

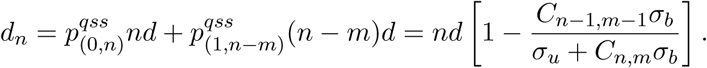

It hence follows that the effective transition rate from group *n* to group *n* + *k* due to a burst of size *k* is given by 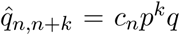 and the effective transition rate from group *n* to group *n* − 1 due to protein decay is given by 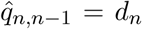. Since we have all effective transition rates of the reduced model, we can now write down the effective reduced master equation in the limit of fast gene switching. Let 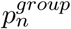 denote the probability of being in group *n*. Then the time-evolution of the reduced model is governed by the reduced master equation:

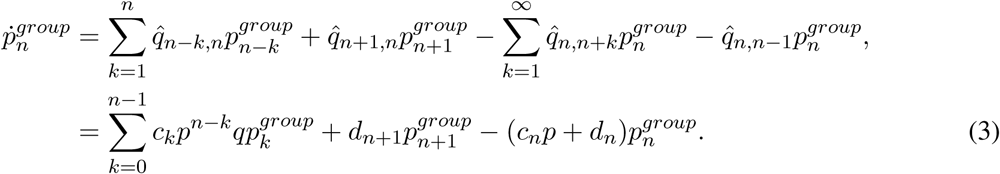

## 3 Solving the reduced master equation in steady-state conditions

### 3.1 General solution

We next solve the reduced master equation exactly in steady-state conditions and thus obtain the steady-state protein number distribution. To solve Eq. (3), we note that it is recursive with respect to the group index *n*. It involves two variables when *n* = 0, three variables when *n* = 1, and so on. Enforcing steady-state conditions by setting the time derivative on the left-hand side to zero, it is straightforward to prove by induction that the solution is given by

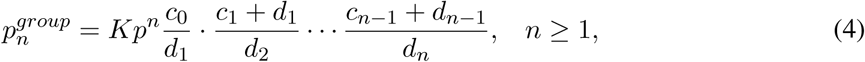

where 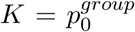 is a normalization constant such that 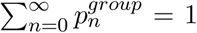. Since *C*_*n,m*_ = *C*_*n*−1,*m*_ + *C*_*n*−1,*m*−1_, it is easy to check that

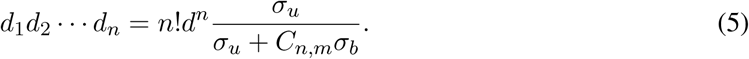

Inserting Eq. (5) into Eq. (4) yields

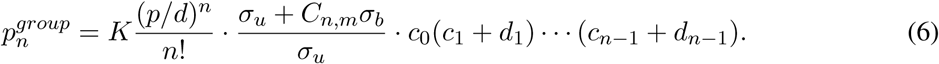

Let *p*_*n*_ = *p*_0,*n*_ + *p*_1,*n*_ denote the probability of having *n* protein copies. Given that there are *n* protein molecules in a single cell, the gene can exist in either microstate (0, *n*) or microstate (1, *n*). Since (0, *n*) is contained in group *n* and (1, *n*) is contained in group *n* + *m*, the probability distribution of protein numbers is given by

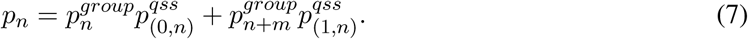

Inserting Eq. (6) into Eq. (7) gives the steady-state distribution of protein numbers:

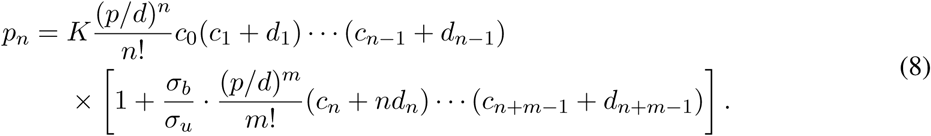

### 3.2 Special cases

We next focus on two important special cases. In the case of *L* = *σ*_*b*_*/σ*_*u*_ ≪ 1, protein binding is much slower than protein unbinding and thus the gene is mostly in the unbound state. In this case, the second term in the square bracket of Eq. (8) is negligible and the effective transcription rate and effective protein decay rate reduce to

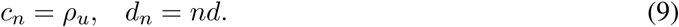

Therefore, the protein number has the negative binomial distribution

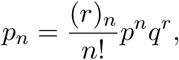

where *r* = *ρ*_*u*_*/d* is the mean number of mRNA copies produced during the protein’s lifetime when the gene is in the unbound state and (*x*)_*n*_ = *x*(*x* + 1) … (*x* + *n* − 1) is the Pochhammer symbol.

Similarly, in the case of *L* = *σ*_*b*_*/σ*_*u*_ ≫ 1, protein binding is much faster than protein unbinding and thus the gene is mostly in the bound state. In this case, the first term in the square bracket of Eq. (8) is negligible and the effective transcription rate and effective protein decay rate reduce to

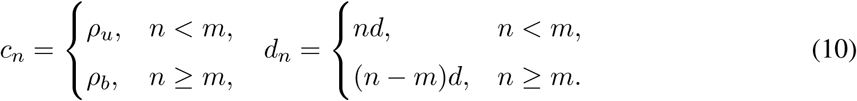

Therefore, the steady-state protein distribution reduces to the negative binomial distribution

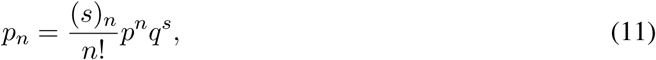

where *s* = *ρ*_*b*_*/d* is the mean number of mRNA copies produced during the protein’s lifetime when the gene is in the bound state.

### 3.3 Testing the accuracy of the analytical solution

As a check of our reduction method and the steady-state analytic solution, we compare it with the numerical solution obtained using FSP [27] for the full master equation given in Eq. (2). The results for positive and negative feedback loops are shown in Fig. 2(a) and Fig. 2(b), respectively. When using FSP, we truncate the state space at a large integer *N* and solve the truncated master equation numerically using the MATLAB function ODE45. The truncation size was chosen as *N* = 5 max(*ρ*_*b*_*B, ρ*_*u*_*B*). Since *ρ*_*b*_*B* and *ρ*_*u*_*B* are the typical protein numbers in the bound and unbound gene states, respectively, the probability that the protein number is outside this truncation size is very small and practically can always be ignored (according to our simulations, *N* = 3 max(*ρ*_*b*_*B, ρ*_*u*_*B*) is already accurate enough). Note that we used FSP rather than the stochastic simulation algorithm (SSA) since in the regime of fast gene switching, the former is much faster computationally than the SSA, because a majority of the time in the SSA is spent simulating gene switching events.

**Fig. 2.**
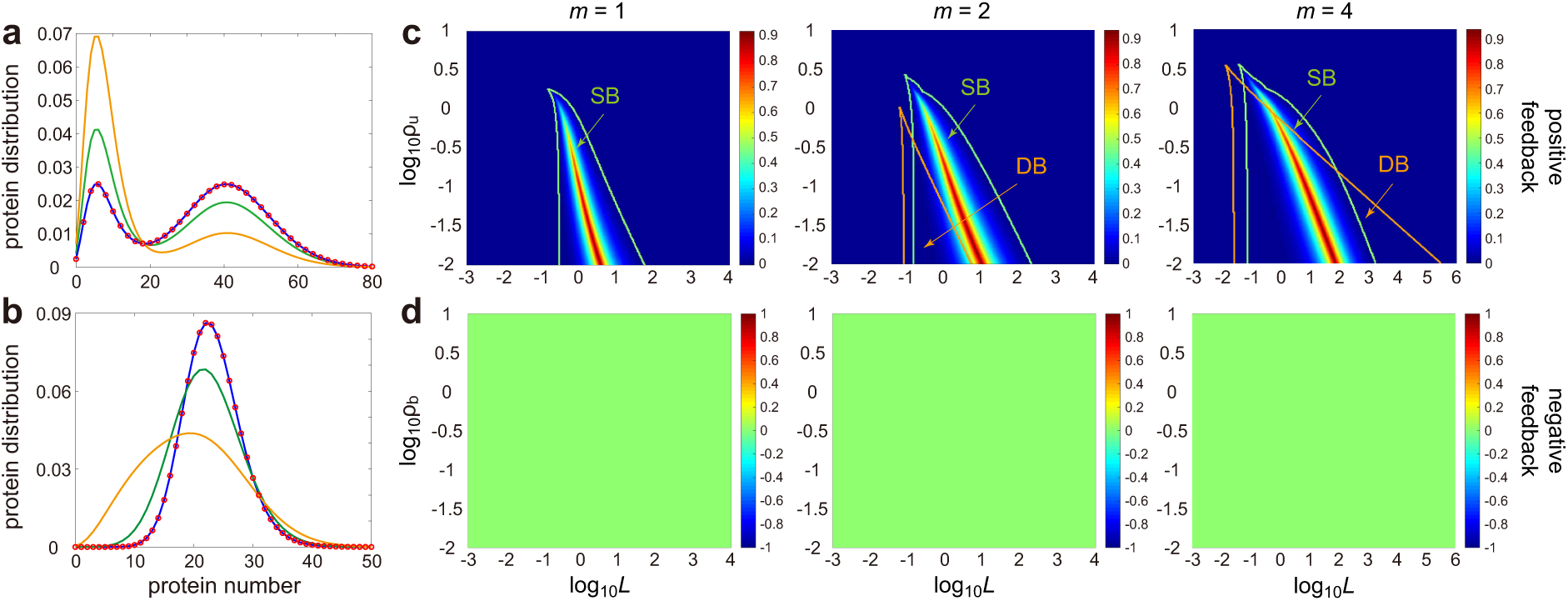
Steady-state behaviour of auto-regulated bursty gene expression with cooperative binding in fast switching conditions. (a),(b) Comparison of the steady-state analytic solution (red circles) with FSP (coloured curves) for different gene switching rates. The model parameters are chosen such that (a) shows a positive feedback loop (*ρ*_*b*_ = 30, *ρ*_*u*_ = 5) and (b) shows a negative feedback loop (*ρ*_*b*_ = 5, *ρ*_*u*_ = 30). The rest of the model parameters in (a),(b) are chosen as *m* = 4, *d* = 1, *L* = 1.16 × 10^*−*4^, *p* = 0.6. The protein unbinding rate is chosen as *σ*_*u*_ = 10^6^ (blue), *σ*_*u*_ = 5, (green) and *σ*_*u*_ = 1 (yellow). (c) Regions in parameter space where SB (stochastic bimodality) and DB (deterministic bistability) are exhibited for a positive feedback loop: the green curve encloses the region for SB and the orange curve for DB. The region designated as SB is that satisfying the criterion *κ >* 0 and the region designated as DB is determined by finding the number of positive real roots of the rate equation given by Eq. (14) in steady-state conditions, *c*(*x*)*B* = *dx*. If this equation has exactly three positive real roots (two stable and one unstable), then the system will show DB. The strength of bimodality *κ* is also shown by the heat map, depicting it as a function of the feedback strength *L*, cooperativity *m* and the transcription rate *ρ*_*u*_ in the unbound gene state under fast gene switching. The model parameters are chosen as *ρ*_*b*_ = 10, *d* = 1, *σ*_*u*_ = 10^6^, *p* = 0.5. (d) Same as (c) but for a negative feedback loop with model parameters chosen as *ρ*_*u*_ = 10, *d* = 1, *σ*_*u*_ = 10^6^, *p* = 0.5. In (c),(d), the cooperativity is chosen as *m* = 1 (left), *m* = 2 (middle) and *m* = 4 (right).

From Fig. 2(a),(b), it is clear that the analytic solution is in excellent agreement with FSP under fast gene switching, but as expected, significant deviations appear for moderate or slow gene switching. Our model predicts both unimodal and bimodal steady-state protein distributions. To gain a deeper insight into bimodal gene expression, we define the strength of bimodality as

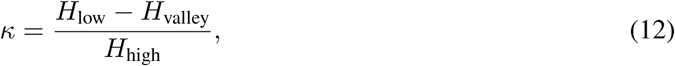

where *H*_low_ and *H*_high_ are the heights of the low and high expression peaks, respectively, and *H*_valley_ is the height of the valley between them. As can be seen from the definition, *κ* is a quantity between 0 and 1 for bimodal protein distributions and is set to 0 for unimodal protein distributions. In general, to display strong bimodality, the following two conditions are necessary: (i) the two peaks should have similar heights and (ii) there should be a deep valley between them. The former ensures that the time periods spent in the low and high expression states are comparable while the latter guarantees that the two expression levels are distinguishable. Clearly, *κ* is large if the two conditions are both satisfied and is small if any one of the two conditions is violated; hence *κ* serves as an effective indicator that characterizes the strength of bimodality [8]. We shall next refer to bimodality in steady-state conditions as stationary bimodality (SB). In Fig. 2(c),(d), we investigate the relationship between SB (characterized by its strength *κ*) and the type of feedback loop, the feedback strength *L* = *σ*_*b*_*/σ*_*u*_, the cooperativity *m* and the smaller one of *ρ*_*b*_ and *ρ*_*u*_ (which represents the transcription rate in the repressed gene state). In the positive feedback case, SB fails to be observed when *ρ*_*u*_ and *ρ*_*b*_ are comparable but can be observed over a wide range of *L* when *ρ*_*u*_ ≪ *ρ*_*b*_ (Fig. 2(c)). In particular, our stochastic model predicts that positive feedback is capable of SB when gene switching is fast even in the absence of cooperative binding (the case of *m* = 1). Increasing cooperativity enlarges the region where SB is observed. On the other hand, a negative feedback loop does not exhibit SB under fast gene switching, independent of whether there is cooperative binding or not (Fig. 2(d)).

Bimodality in the stochastic model can be seen as arising due to switching between two effective phenotypic states of the system. A different definition of switching behaviour, albeit the classical one, is that stemming from deterministic rate equations: if the steady-state solution of these equations has two stable fixed points then the system is bistable and if it has only one stable fixed point then it is monostable. We shall refer to the former as deterministic bistability (DB) to clearly distinguish it from SB. The deterministic rate equations follow from a mean-field approximation of the full CME of the auto-regulatory feedback loop and is given by the following set of ODEs (see Appendix A for its derivation):

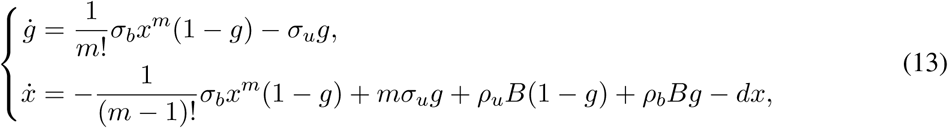

where *g* is the mean number of genes in the bound state and *x* = ⟨*n*⟩ is the mean protein number. In the limit of fast gene switching, the gene component is in fast equilibrium and thus we have

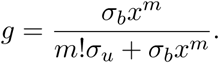

Inserting this equation into the equation for mean protein number, we obtain the following effective rate equation:

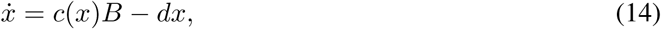

where

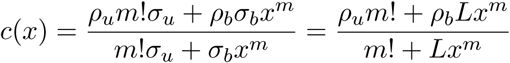

is the effective transcription rate.

In Fig. 2(c),(d), we also depict the regions of parameter space where DB is observed. For negative feedback loops, no DB is exhibited, as was also the case for SB (Fig. 2(d)). However for positive feedback loops, the regions of SB and DB are different (Fig. 2(c)). For no cooperativity (*m* = 1), SB is observed but not DB; for moderate cooperativity (*m* = 2), the region of DB becomes significantly enlarged though still much smaller than that of SB; for high cooperativity (*m* = 4), the regions of SB and DB overlap to a considerable extent. Hence the differences between SB and DB are most apparent for positive feedback loops with low cooperativity. Note that within the mean-field approximation, DB is associated with a bimodal distribution where each mode corresponds to one of the stable states. Hence in the regions where there is SB but not DB, we can say that noise induces the bimodality of the distribution. In contrast, in the regions where there is DB but not SB, we can say that noise induces the unimodality of the distribution.

## 4 Solving the time-dependent reduced master equation

### 4.1 Analytic solution

Following the method proposed in [28], we next solve the reduced master equation in time and thus obtain the time-dependent protein number distribution. For convenience, we truncate the state space at a large integer *N*. Let 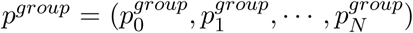 denote the time-dependent solution of the reduced model. It then follows that the reduced master equation given by Eq. (3) can be rewritten in matrix form as

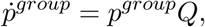

where

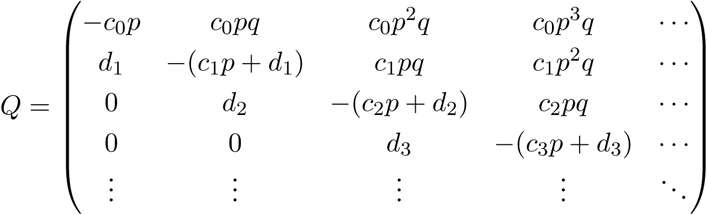

is the generator matrix of the reduced model. The solution to this ODE can be represented using the matrix exponential as

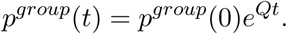

It is usually difficult to compute the matrix exponential analytically. In general, we need to calculate the eigenvalues and eigenvectors of *Q*. However, the special structure of *Q* allows us to bypass the eigenvector calculation. By Cauchy’s integral formula for matrices [29], for any continuous function *f*, we have

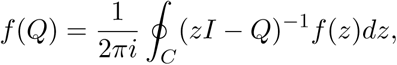

where *C* is an arbitrary simple closed curve in the complex plane that contains all eigenvalues of *Q* in its interior. If we take *f* (*z*) = *p*^*group*^(0)*e*^*tz*^, then we obtain

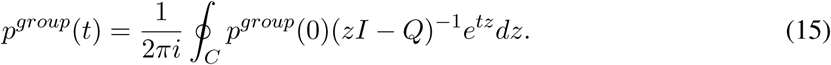

Suppose now that initially we start from a fixed group *n*_0_; then the initial distribution of the reduced model is given by 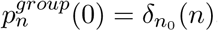, where 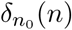 is a Kronecker delta that takes the value of 1 when *n* = *n*_0_ and the value of 0 otherwise. We shall explain later how to extend our results to more general initial distributions. Under the point initial distribution, Eq. (15) can be simplified to

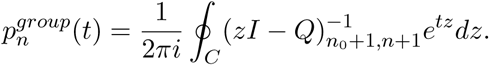

By Cramer’s rule of computing the inverse matrix, we can prove that

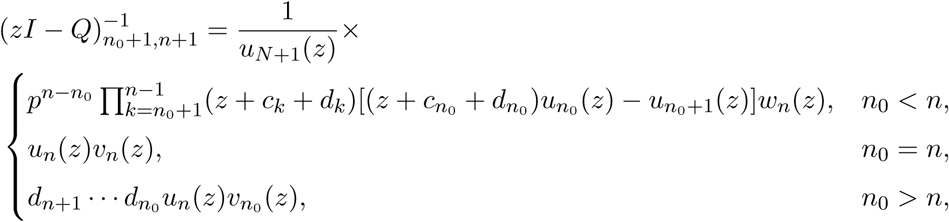

where *u*_*n*_(*z*) are polynomials defined recursively by

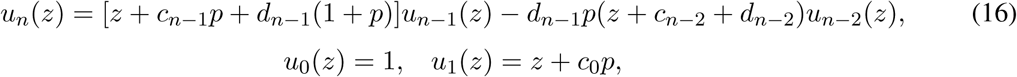

with *u*_*N*+1_(*z*) = det(*zI* − *Q*) being the characteristic polynomial of *Q*, and *v*_*n*_(*z*) are polynomials defined recursively by

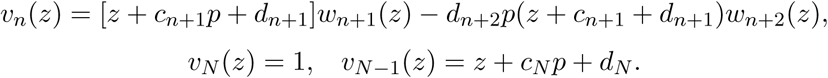

Here *v*_*n*_(*z*) are defined with the aid of another sequence of polynomials *w*_*n*_(*z*), which are defined recursively by

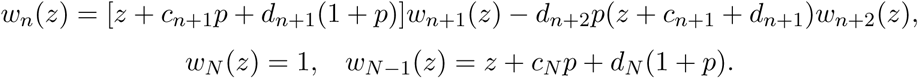

Therefore, we obtain

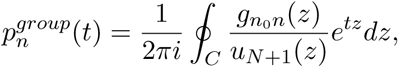

where 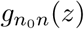 are polynomials defined as

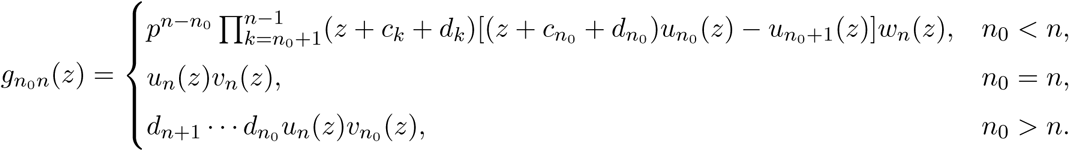

Since *u*_*N*+1_(*z*) is the characteristic polynomial of *Q*, we have

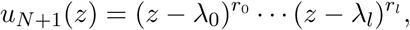

where *λ*_0_, …, *λ*_*l*_ are all pairwise distinct eigenvalues of *Q* with *r*_0_, …, *r*_*l*_ being their multiplicities, respectively. We can then apply Cauchy’s residue theorem:

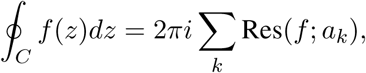

where *a*_*k*_ are all singularities of *f* inside the simple closed curve *C*. In our current case, the singularities are all the eigenvalues *λ*_*k*_ and thus

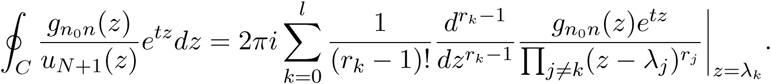

Therefore, once we have known all eigenvalues of *Q*, the time-dependent solution of the reduced model is given by

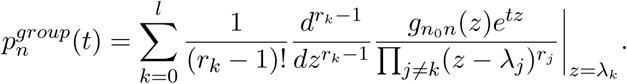

In most cases, the eigenvalues of *Q* are mutually different (any matrix can be approximated by such matrices to any degree of accuracy). In this case, we have *l* = *N* and *r*_0_ = … = *r*_*l*_ = 1 and thus the time-dependent solution can be simplified as

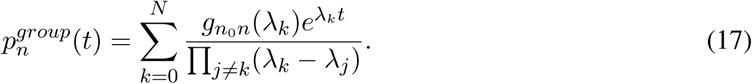

Since we have found the transient solution of the reduced model, inserting Eq. (17) into Eq. (7) finally gives the time-dependent distribution of protein numbers:

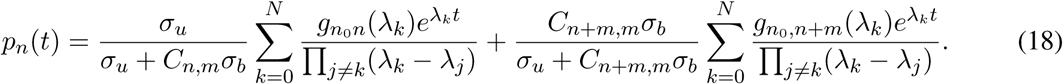

In the general case that the reduced model starts from a general initial distribution *p*_*n*_(0) = *π*_*n*_, the transition solution is given by

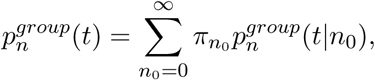

where 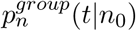 is the transient solution given that the reduced model starts from group *n*_0_. In what follows, we assume that the initial protein number is zero, i.e. *n*_0_ = 0, and the gene is initially in the unbound state, unless otherwise stated. This assumption is common in studies comparing experimental data with deterministic model predictions for the time dependence of autoregulation [30]. We emphasize here that although our theory is presented in the framework of auto-regulated bursty gene expression, the time-dependent analytic solution derived above can be applied to an arbitrary bursty birth-death processes with general birth rate *c*_*n*_ and general death rate *d*_*n*_. This makes our method widely applicable beyond the framework of stochastic gene expression.

### 4.2 Convergence to the steady-state solution

In fact, the steady-state solution obtained earlier can be recovered from the time-dependent solution by taking *t* → ∞. By the Perron-Frobenius theorem [31], when the system is ergodic, the generator matrix *Q* must have zero as its eigenvalue with multiplicity one and all other eigenvalues must have negative real parts:

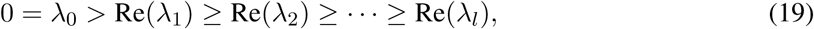

where Re(*z*) is the real part of *z*. Therefore, the only term in Eq. (17) independent of time *t* is the first term and all other terms tend to zero exponentially fast as *t* → ∞. Since the system is ergodic, the steady-state solution is independent of the choice of the initial distribution. As a result, we can take *n*_0_ to be sufficiently large, allowing us to focus on to the case of *n*_0_ *> n*, which is now

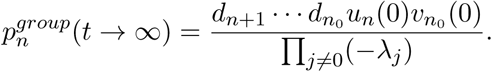

Since the term 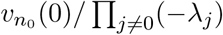 contributes the same to each term, we can treat it as a normalizing factor. We can also do the same with 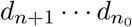, which contributes the same as (*d*_1_ … *d*_*n*_)^−1^ up to normalization. Therefore, the steady-state solution has the following simplified expression:

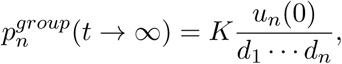

where *K* is a normalization constant. Moreover, we can prove by induction that

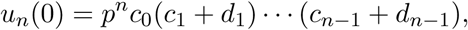

which finally leads to

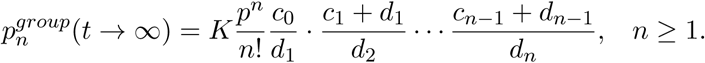

This is exactly the same as Eq. (4) obtained in the previous section.

### 4.3 Approximate eigenvalues for the cases of fast and slow protein binding

Our analytic solution for the time-dependent protein distribution as given by Eq. (18) depends on all eigenvalues of the generator matrix *Q*. In general, it is very difficult to compute these eigenvalues analytically. However, as we now show, this can be done for two special cases, namely when protein binding is much slower or much faster than protein unbinding.

We first focus on the case of *L* ≪ 1, i.e. protein binding is much slower than protein unbinding. In this case, all the eigenvalues of *Q* are approximately given by (see Appendix B for details)

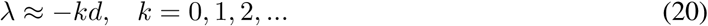

In other words, when *L* ≪ 1, all the approximate eigenvalues of *Q* are nonpositive integer multiples of the protein decay rate.

We next focus on the case of *L* ≫ 1, i.e. protein binding is much faster than protein unbinding. In this case, all the eigenvalues of *Q* are approximately given by (see Appendix C for details)

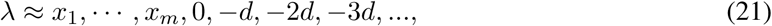

where *x*_1_, …, *x*_*m*_ are all the zeros of the polynomial *u*_*m*_(*z*) of degree *m* defined in Eq. (16). Using the approximation given in Eq. (10), the polynomial *u*_*m*_(*z*) is defined recursively by

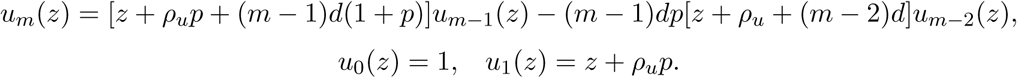

In other words, when *L* ≫ 1, all the approximate eigenvalues of *Q* are nonpositive integer multiples of the protein decay rate combined with all the zeros of the polynomial *u*_*m*_(*z*). In particular, in the non-cooperative case of *m* = 1, we have *x*_1_ = −*ρ*_*u*_*p* and thus all the approximate eigenvalues of *Q* are given by

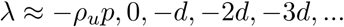

In the cooperative case of *m* = 2, we have

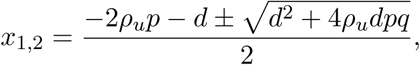

and thus all the approximate eigenvalues of *Q* are given by

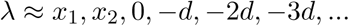

Substituting these approximate eigenvalues in Eq. (18) gives the approximate time-dependent protein distribution.

To verify the accuracy of the approximate time-dependent solution, we compare it with FSP in the regimes of *L* ≪ 1 (Fig. 3(a)) and *L* ≫ 1 (Fig. 3(b)). Clearly, the approximate solution is in excellent agreement with FSP. With the parameters chosen in Fig. 3, the first eight real and approximate eigenvalues of *Q* are listed in Table 1, from which we can see that the approximate eigenvalues are very accurate in the two limiting regimes. When *L* is neither too large nor too small, we can compute the eigenvalues of *Q* numerically and substituting them in Eq. (18) to obtain the semi-exact time-dependent protein distribution. The transient solution obtained in this way is in full agreement with FSP for a large range of model parameters and over time in the regime of fast gene switching (Fig. 4).

**Table 1.**
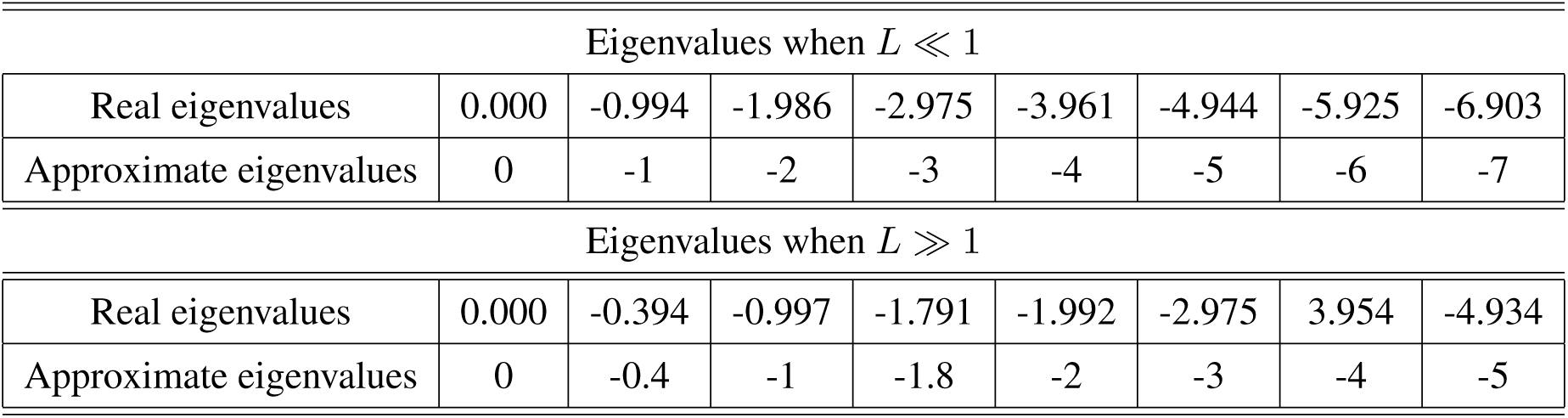
The first eights real and approximate eigenvalues of the generator matrix *Q* when protein binding is much slower (*L* ≪ 1) or much faster (*L* ≪ 1) than protein unbinding. In the case of *L* ≪ 1, the model parameters are chosen as *m* = 2, *ρ*_*b*_ = 10, *ρ*_*u*_ = 5, *d* = 1, *σ*_*u*_ = 10^6^, *L* = 10^*−*4^, *p* = 0.6 and the approximate eigenvalues are computed using Eq. (20). In the case of *L* ≪ 1, the model parameters are chosen as *m* = 2, *ρ*_*b*_ = 10, *ρ*_*u*_ = 1, *d* = 1, *σ*_*u*_ = 10^4^, *L* = 10, *p* = 0.6 and the approximate eigenvalues are computed using Eq. (21). Note that these two sets of parameters (and the associated eigenvalues) correspond to those used to calculate the time-dependent distributions in Fig. 3(a),(b).

| Eigenvalues when $L \ll 1$ | | | | | | | | |
| --- | --- | --- | --- | --- | --- | --- | --- | --- |
| Real eigenvalues | 0.000 | -0.994 | -1.986 | -2.975 | -3.961 | -4.944 | -5.925 | -6.903 |
| Approximate eigenvalues | 0 | -1 | -2 | -3 | -4 | -5 | -6 | -7 |

Table 1. **The first eight real and approximate eigenvalues of the generator matrix $Q$ when protein binding is much slower ( $L \ll 1$ ) or much faster ( $L \gg 1$ ) than protein unbinding.**
| Eigenvalues when $L \gg 1$ | | | | | | | | |
| --- | --- | --- | --- | --- | --- | --- | --- | --- |
| Real eigenvalues | 0.000 | -0.394 | -0.997 | -1.791 | -1.992 | -2.975 | 3.954 | -4.934 |
| Approximate eigenvalues | 0 | -0.4 | -1 | -1.8 | -2 | -3 | -4 | -5 |

**Fig. 3.**
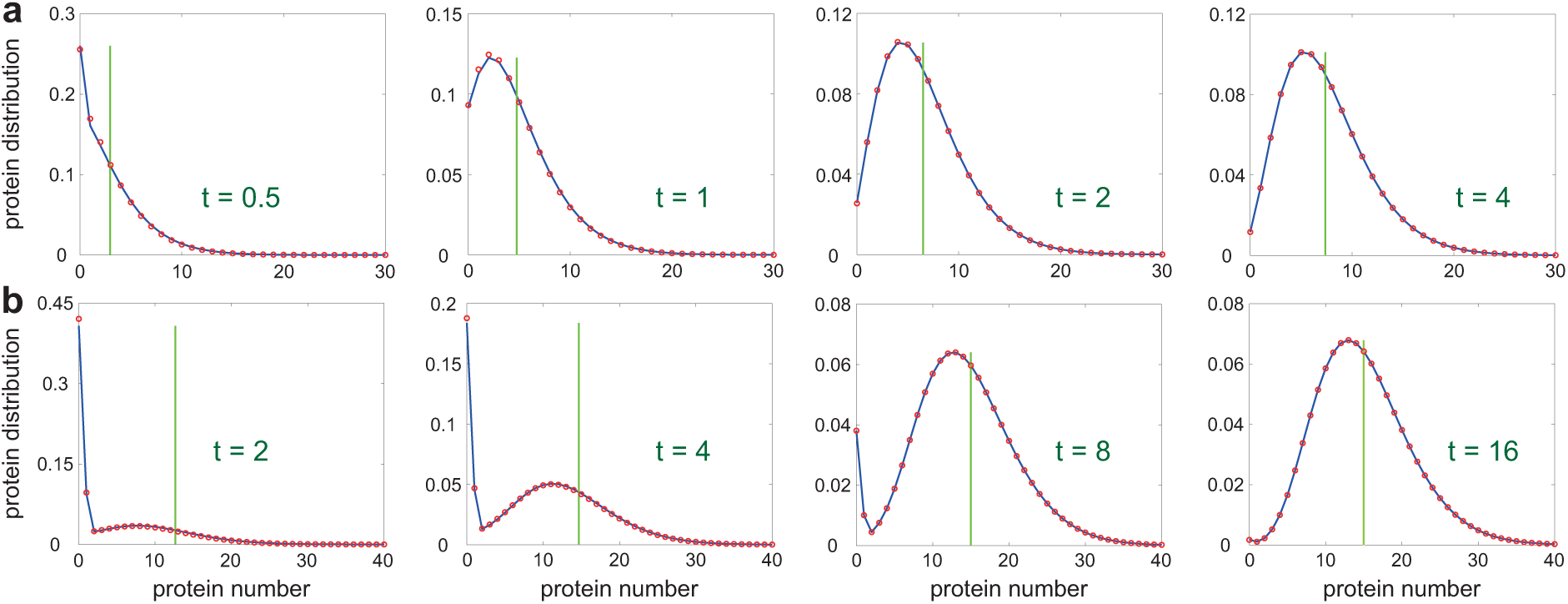
Comparison of the approximate time-dependent solution (red circles) with FSP (blue curve) when protein binding is much slower (*L* ≪ 1) or much faster (*L* ≫ 1) than protein unbinding. (a) Case of slow protein binding. The model parameters are chosen as *m* = 2, *ρ*_*b*_ = 10, *ρ*_*u*_ = 5, *d* = 1, *σ*_*u*_ = 10^6^, *L* = 10^*−*4^, *p* = 0.6. The red circles are obtained by substituting the approximate eigenvalues given by Eq. (20) in our analytic solution given by Eq. (18) for *N* = 200. (b) Case of fast protein binding. The model parameters are chosen as *m* = 2, *ρ*_*b*_ = 10, *ρ*_*u*_ = 1, *d* = 1, *σ*_*u*_ = 10^4^, *L* = 10, *p* = 0.6. The red circles are obtained by substituting the approximate eigenvalues given by Eq. (21) in our analytic solution given by Eq. (18) for *N* = 200. The green vertical line in (a) and (b) shows the mean protein number predicted by the deterministic rate equation.

**Fig. 4.**
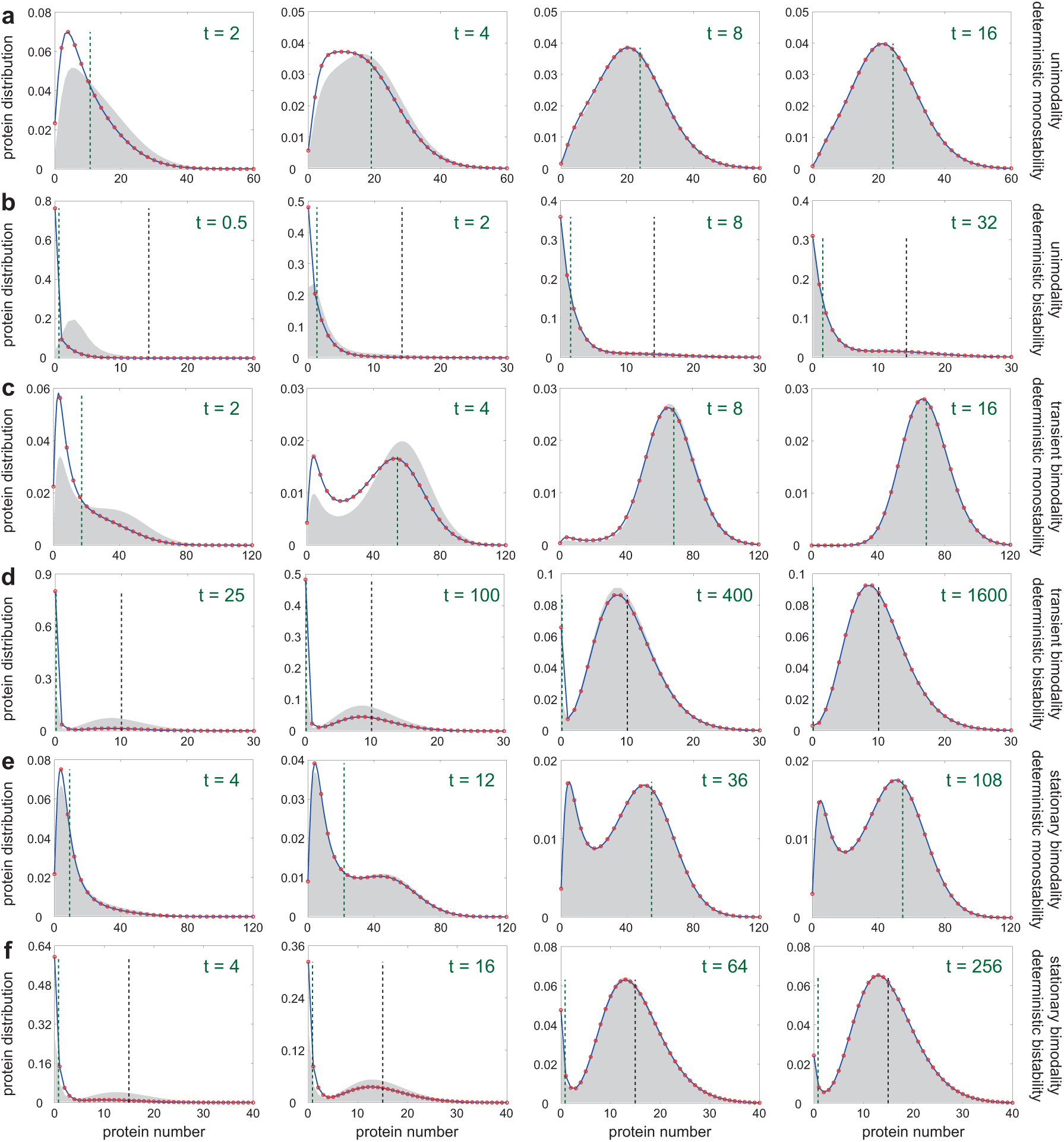
Comparison of the semi-analytical time-dependent solution with FSP for six distinct types of dynamic behaviours at four time points. For an initial protein number equal to zero and the gene is initially in the unbound state, the analytical solution is shown by red circles while FSP is shown by the solid blue line. Specifically the red circles are obtained by substituting the numerically found eigenvalues of the generator matrix *Q* (truncated at *N* = 200) in Eq. (18). The dash green vertical line shows the mean protein number predicted by the deterministic rate equations. The black vertical line shows the other stable fixed point (if there exists one) of the deterministic rate equation. For an initial protein distribution given by a Poisson with mean 5, we show the numerical time-dependent distribution obtained using FSP in solid grey. U, TB and SB refer to the following three properties: unimodality at all times, transient bimodality at intermediate times (unimodal at short and long times) and bimodality at long times (unimodal at short times), respectively. DM and DB refer to deterministic monostability and bistability, respectively. (a) U+DM. The model parameters are chosen as *m* = 2, *ρ*_*b*_ = 20, *ρ*_*u*_ = 5, *d* = 1, *σ*_*u*_ = 10^6^, *σ*_*b*_ = 0.01*σ*_*u*_, *p* = 0.6. (b) U+DB. The model parameters are chosen as *m* = 4, *ρ*_*b*_ = 10, *ρ*_*u*_ = 1, *d* = 1, *σ*_*u*_ = 10^6^, *σ*_*b*_ = 0.009*σ*_*u*_, *p* = 0.6. (c) TB+DM. The model parameters are chosen as *m* = 2, *ρ*_*b*_ = 50, *ρ*_*u*_ = 5, *d* = 1, *σ*_*u*_ = 10^6^, *σ*_*b*_ = 0.0045*σ*_*u*_, *p* = 0.6. (d) TB+DB. The model parameters are chosen as *m* = 4, *ρ*_*b*_ = 10, *ρ*_*u*_ = 0.1, *d* = 1, *σ*_*u*_ = 10^6^, *σ*_*b*_ = 250*σ*_*u*_, *p* = 0.5. (e) SB+DM. The model parameters are chosen as *m* = 2, *ρ*_*b*_ = 50, *ρ*_*u*_ = 4, *d* = 1, *σ*_*u*_ = 10^6^, *σ*_*b*_ = 0.0016*σ*_*u*_, *p* = 0.6. (f) SB+DB. The model parameters are chosen as *m* = 4, *ρ*_*b*_ = 10, *ρ*_*u*_ = 5, *d* = 0.5, *σ*_*u*_ = 10^6^, *σ*_*b*_ = 0.3*σ*_*u*_, *p* = 0.6.

## 5 Classification of the time trajectories of an auto-regulating gene

### 5.1 Dynamical phase diagrams

According to both the numerical solution of the full CME and the semi-analytical solution shown in Fig. 4, our auto-regulatory gene expression model can exhibit three different types of dynamic behaviours: (i) the protein distribution is unimodal at all times (Fig. 4(a),(b)), (ii) the protein distribution is unimodal at small and large times and is bimodal at intermediate times (Fig. 4(c),(d)), and (iii) the protein distribution is unimodal at small times and is bimodal at large times (Fig. 4(e),(f)). To distinguish between them, we refer to (i) as unimodality (U), to (ii) as transient bimodality (TB), and to (iii) as stationary bimodality (SB; this has already been introduced earlier).

Each type of dynamic behaviour can be further divided into two phases according to whether the deterministic model shows monostability (DM) or bistability (DB). Hence the dynamic behaviour of our auto-regulatory gene expression model can be classified into six possible phases (Table 2). Phases 1 and 6 (U+DM and SB+DB, respectively) are cases where the stochastic and deterministic models both predict the same behaviour; in other words the presence of noise has no effect on determining the number of modes of the protein distribution. In contrast, in the other four phases 2-5, noise plays an important role in the creation or destruction of bimodality. We further note that TB (the behaviour of the stochastic model in phases 3 and 4) is a purely stochastic effect since bistability in the deterministic model can only be determined at the steady state. The numerical solution obtained using FSP and the analytical solution indicate that each of the six dynamical phases can appear when model parameters are appropriately chosen (Fig. 4(a)-(f)). Note that the existence of these phases is not specific to an initial condition given by a delta function; as shown by the grey distributions in Fig. 4, the same phases are also found when the initial condition follows a Poisson distribution. To determine the regions for the six phases in parameter space, we illustrate the *L*-*ρ*_*u*_ and *L*-*B* phase diagrams for positive feedback loops (Fig. 5(a),(b)) and the *L*-*ρ*_*b*_ and *L*-*B* phase diagrams for negative feedback loops (Fig. 5(c),(d)).

**Table 2.**
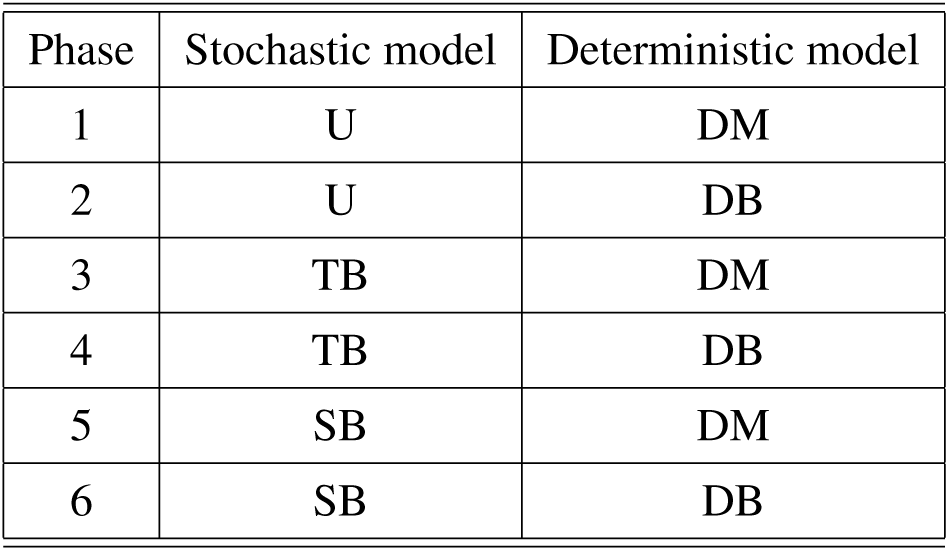
Six dynamical phases of auto-regulated bursty gene expression in fast switching conditions. There are three possible phases for the stochastic model (U, TB, and SB) and two possible phases for the deterministic model (DM and DB), so that in sum there are six possible dynamical phases.

**Fig. 5.**
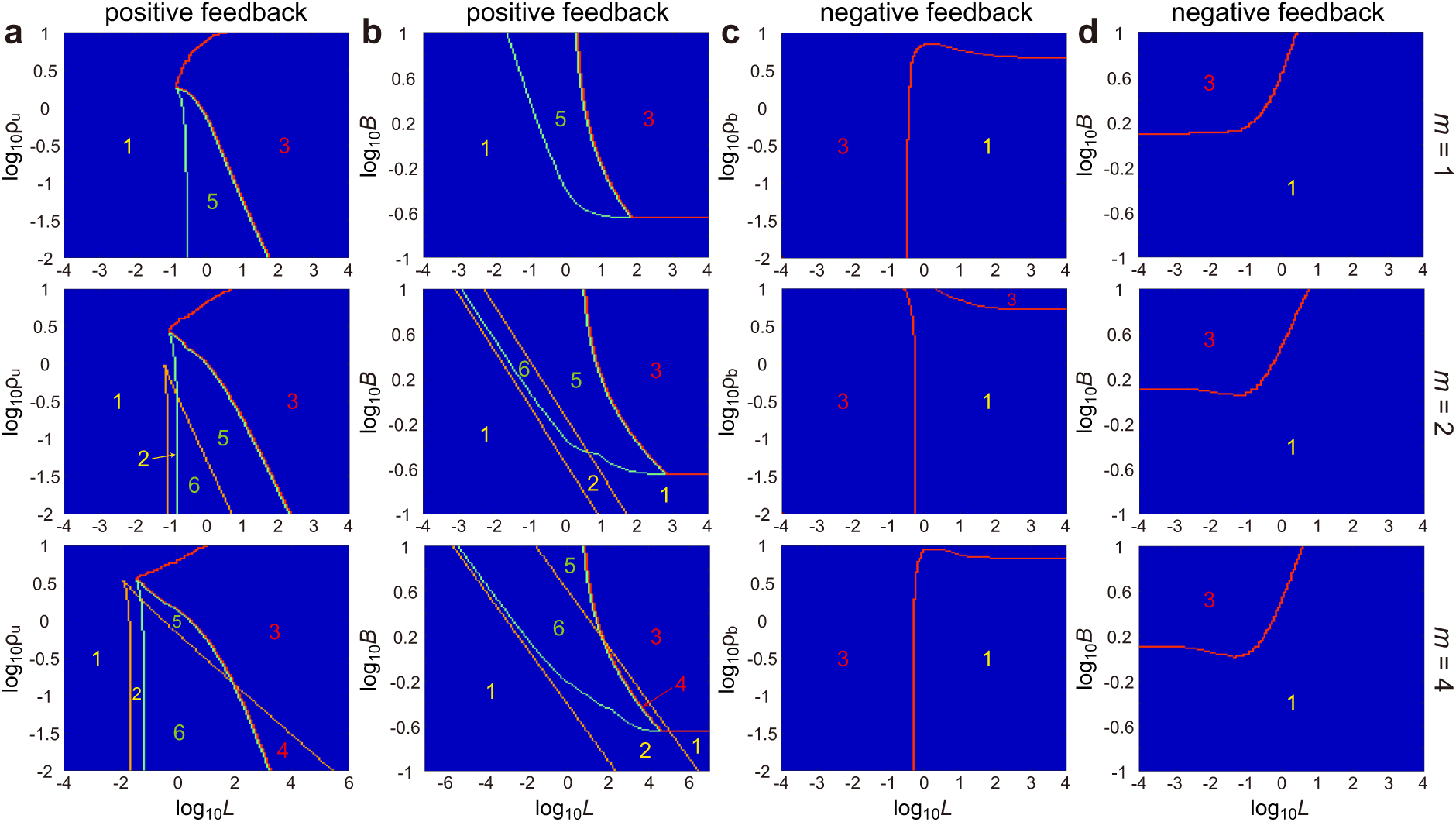
Dynamical phase diagrams for auto-regulated bursty gene expression with cooperative binding in fast switching conditions. Different phases are labelled according to the classification in Table 2. (a) *L*-*ρ*_*u*_ phase diagram for positive feedback loops. The model parameters are chosen as *ρ*_*b*_ = 10, *d* = 1, *σ*_*u*_ = 10^6^, *B* = 1. (b) *L*-*B* phase diagram for positive feedback loops. The model parameters are chosen as *ρ*_*b*_ = 10, *ρ*_*u*_ = 0.1, *d* = 1, *σ*_*u*_ = 10^6^. (c) *L*-*ρ*_*b*_ phase diagram for negative feedback loops. The model parameters are chosen as *ρ*_*u*_ = 10, *d* = 1, *σ*_*u*_ = 10^6^, *B* = 2. (b) *L*-*B* phase diagram for negative feedback loops. The model parameters are chosen as *ρ*_*b*_ = 0.1, *ρ*_*u*_ = 10, *d* = 1, *σ*_*u*_ = 10^6^. In each phase diagram, we keep *σ*_*u*_ as a constant and change the value of *L* by tuning *σ*_*b*_. In (a)-(d), the cooperativity is chosen as *m* = 1 (up), *m* = 2 (middle) and *m* = 4 (down).

We first focus on the positive feedback case (Fig. 5(a),(b)). From the phase diagrams, it can be seen that the system exhibits TB when the feedback strength *L* is large. The dependence of TB on the mean protein burst size *B* is less strong though it appears that *B* must be sufficiently large too (in fact, *B* must be greater than 2*/*(*s* − 1), as will be proved later). In most biologically relevant cases, the transcription rates in the two gene states are not of the same order of magnitude, i.e. *ρ*_*u*_ ≪ *ρ*_*b*_, and the mean burst size *B* is large. In this situation, as the feedback strength *L* increases, a positive auto-regulatory gene network typically undergoes two successive stochastic bifurcations, from the U phase to the SB phase and then to the TB phase (Fig. 5(a),(b)). In both phase diagrams, there is a triple point separating the U, SB, and TB phases of the stochastic model — *this is analogous to the triple point in the phase transition between solid, liquid, and gaseous states of a substance due to the effects of temperature and pressure* [32]. Note that all the six phases appear for high cooperativity (*m* = 4), but for no cooperativity (*m* = 1) only three of the six phases are observed; hence increasing nonlinearity in the mass action law for binding kinetics increases the richness of the system’s temporal behaviour. The rarest phases are phases 2 and 4 (U+DB and TB+DB, respectively), implying that when cooperativity is sufficiently high such that there is deterministic bistability, the stochastic model is most likely to exhibit stationary bimodality, i.e. to be in the phase 6 (SB+DB).

We next focus on the negative feedback case (Fig. 5(c),(d)). Since the stochastic model cannot produce SB and the deterministic model cannot produce DB (as determined in Section 3), a negative feedback loop only has two types of dynamic behaviours, i.e. phases 1 and 3 (U+DM and TB+DM, respectively). This is in contrast to a positive feedback loop which possesses all six dynamical phases. From the phase diagrams, it can be seen that the system exhibits TB when the feedback strength *L* is relatively small but the mean protein burst size *B* is relatively large. For negative feedback loops, the value of the transcriptional rate *ρ*_*u*_ in the repressed gene state practically seems uncorrelated with TB. In the typical case that the transcription rates in the two gene states are not of the same order of magnitude, i.e. *ρ*_*b*_ ≪ *ρ*_*u*_, and the mean burst size *B* is large, as the feedback strength *L* increases, a negative auto-regulatory gene network undergoes a stochastic bifurcation from the TB phase to the U phase (Fig. 5(c),(d)).

In both positive and negative feedback cases, we find that an increased protein burst size broadens the region of TB to a large extent. When all model parameters are fixed (except a possible interchange between *ρ*_*u*_ and *ρ*_*b*_), a negative feedback loop requires a significantly larger burst size to produce TB than a positive feedback loop (Fig. 5(b),(d)).

### 5.2 Analytical estimation of the observation time window for transient bimodality

Phases displaying TB are the most interesting since they have no deterministic counterpart. Now we shall use the theory developed earlier in Section 4.1 to verify the existence of TB theoretically and more importantly to estimate the observation time window over which it can be detected. For convenience, we rewrite Eq. (18) as

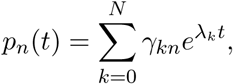

where

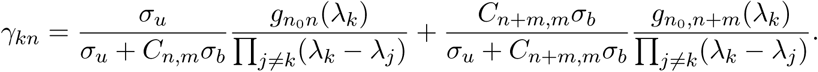

The coefficients associated with the exponential functions satisfy (see Appendix D for details)

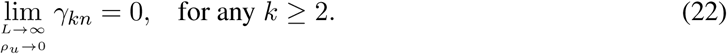

This shows that when *L* ≫ 1 (fast protein binding) and *ρ*_*u*_ ≪ *ρ*_*b*_ (the transcription rate in the repressed gene state is much smaller than that in the active gene state), all terms can be ignored except for the first two exponential terms:

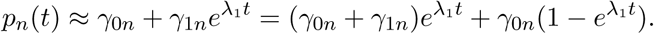

Since the the initial protein number is assumed to be zero, we have

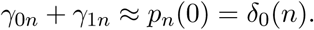

Moreover, since the system is ergodic and *L* ≫ 1, it follows from Eq. (11) that

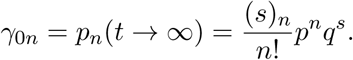

Therefore, at each intermediate time *t*, the protein number has a zero-inflated negative binomial (ZINB) distribution

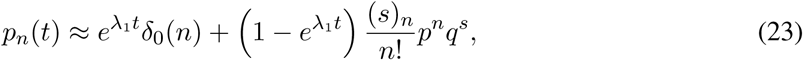

which is a mixture of a point mass at zero and a negative binomial distribution with their coefficients depending on time *t*. When *t* is relatively large, the time-dependent protein distribution only depends on the first few exponential terms (the steady-state protein distribution only depends on the first exponential term), while when *t* is very small, the time-dependent protein distribution depends on all exponential terms. Since we only retain the first two exponential terms in the approximation, the ZINB distribution may deviate from the real distribution when *t* is extremely small.

To understand when bimodality manifests, we note that the mode of the negative binomial part is given by

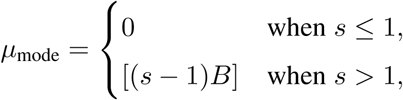

where [*x*] denotes the integer part of *x*. Hence the ZINB distribution peaks at both zero and the non-zero mode [(*s* − 1)*B*], if and only if *p*_0_(*t*) *> p*_1_(*t*) and *µ*_mode_ ≥ 2, i.e.

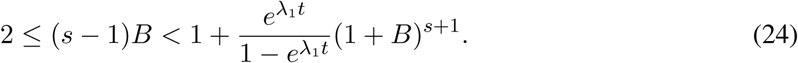

From Eq. (24), we see that there is a critical time

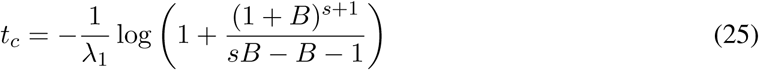

such that bimodality occurs if and only if (*s* − 1)*B* ≥ 2 and 0 *< t < t*_*c*_ (Fig. 6). Note that while the ZINB distribution predicts bimodality at very small times, the real distribution may be unimodal because the approximate distribution may deviate from the real one when *t* ≪ 1 (Fig. 6). Nevertheless the calculation provides an accurate estimate of the observation time window for TB and furthermore confirms the observation in Fig. 5 that for a positive feedback loop, the phenomenon occurs when the burstiness of protein expression is sufficiently large and protein binding is much faster than unbinding.

**Fig. 6.**
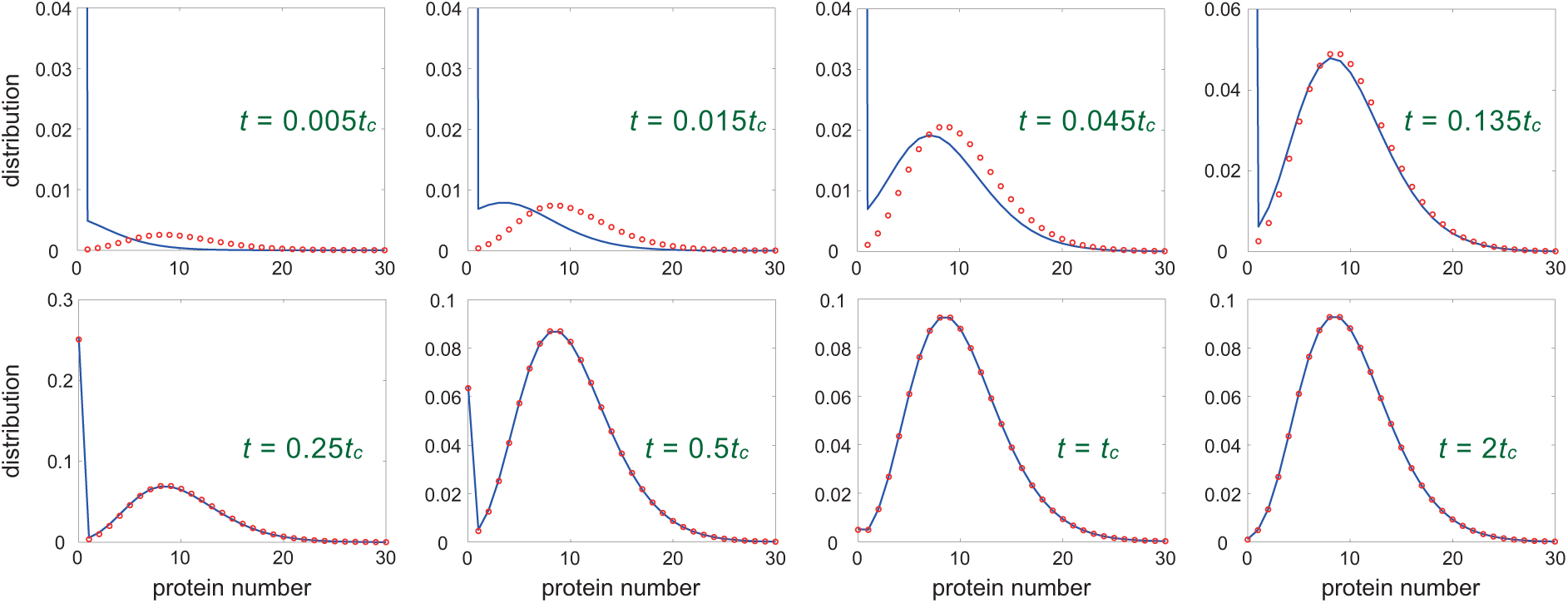
Transient bimodality. When *ρ*_*u*_ ≪ *ρ*_*b*_ and *B* ≥ 2*/*(*s* − 1), a positive feedback loop produces TB when *L* ≫ 1. The blue curve shows the real time-dependent protein distribution simulated using FSP and the red circles show the approximate solution given by Eq. (23). The model parameters are chosen as *m* = 1, *ρ*_*b*_ = 10, *ρ*_*u*_ = 0.1, *σ*_*u*_ = 10^3^, *L* = 100, *p* = 0.5. The critical time for TB is computed using Eq. (25) with *λ*^1^ ≈ − *ρ*_*u*_*p* (the approximate eigenvalues for *m* = 1 and *L* ≫ 1 are computed analytically in Appendix C). Note that FSP confirms the theoretical prediction that TB disappears when *t > t*_*c*_.

Here we have looked at TB for positive feedback. As we have shown in Fig. 5, the phenomenon also exists for negative feedback when protein binding is slow compared to unbinding. Unfortunately the time-dependent solution in this case depends on many exponential terms (often *>* 10 terms) which makes it almost impossible to analytically estimate the observation time window using the same method as we have carried out for positive feedback.

### 5.3 Relationship between the phases, the relaxation time and the protein mean

From our numerical simulations using FSP in Fig. 4, we observe that the relaxation time of our auto-regulatory gene expression model is closely related to its dynamic behaviour. When the system shows U, it relaxes to the steady state rapidly (Fig. 4(a),(b)). When the system shows SB, it takes a much longer time to reach the steady state (Fig. 4(e),(f)). However, when the system shows TB, it can either relax rapidly Fig. 4(c) or relax very slowly Fig. 4(d).

To further study the possible link between the dynamical phases of the system and the relaxation time, we compute the relaxation rate (the inverse of the relaxation time) of our stochastic model across large regions of parameter space. Note that here the relaxation rate is to a good approximation given by *γ* = |Re(*λ*_1_)|, i.e. the spectral gap between the zero eigenvalue (which is associated with the steady state) and the first nonzero eigenvalue (which is associated with the slowest transient decay) of the generator matrix *Q* [33]. In Fig. 7, using heat maps, we show the size of the relaxation rate as a function of the parameters *L, ρ*_*u*_, *ρ*_*b*_ and *B* for positive feedback loops (Fig. 7(a),(b)) and for negative feedback loops (Fig. 7(c),(d)). The regions of parameter space where each of the three phases (U, SB and TB) manifests are also shown.

**Fig. 7.**
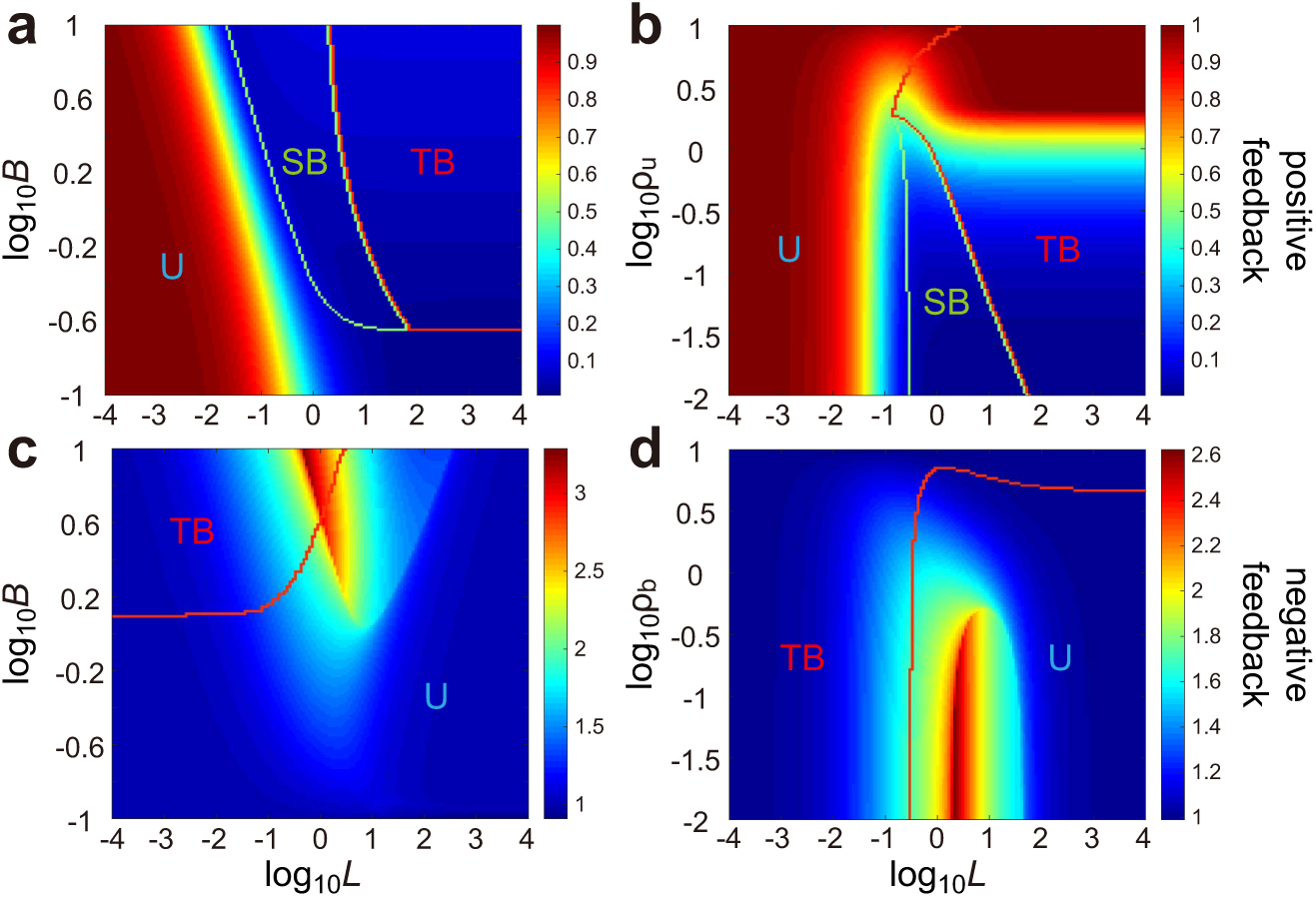
Relaxation kinetics for auto-regulated bursty gene expression. (a) Heat plot shows the relaxation rate *γ* as a function of the feedback strength *L* and the mean protein burst size *B* for positive feedback loops. The model parameters are chosen as *m* = 1, *ρ*_*b*_ = 10, *ρ*_*u*_ = 0.1, *d* = 1, *σ*_*u*_ = 10^6^. (b) Heat plot shows the relaxation rate *γ* as a function of the feedback strength *L* and the transcription rate *ρ*_*u*_ in the unbound gene state for positive feedback loops. The model parameters are chosen as *m* = 1, *ρ*_*b*_ = 10, *d* = 1, *σ*_*u*_ = 10^6^, *B* = 1. (c) Heat plot shows the relaxation rate *γ* as a function of the feedback strength *L* and the mean protein burst size *B* for negative feedback loops. The model parameters are chosen as *m* = 1, *ρ*_*b*_ = 0.1, *ρ*_*u*_ = 10, *d* = 1, *σ*_*u*_ = 10^6^. (d) Heat plot shows the relaxation rate *γ* as a function of the feedback strength *L* and the transcription rate *ρ*_*b*_ in the bound gene state for negative feedback loops. The model parameters are chosen as *m* = 1, *ρ*_*u*_ = 10, *d* = 1, *σ*_*u*_ = 10^6^, *B* = 2. Different phases (U, TB, and SB) are marked on each figure to show the relationship between the relaxation rate and the dynamical phases.

In the positive feedback case, we find that SB (phases 5 and 6) is always contained in the subregion with a small relaxation rate. This clearly shows that SB significantly prolongs the relaxation time of stochastic gene expression. In contrast, U (phases 1 and 2) is associated with a large relaxation rate and hence a short relaxation time. Moreover, we find that while TB (phases 3 and 4) occupies a large portion of the subregion with a small relaxation rate, it also occupies a substantial portion of the subregion with a large relaxation rate. This shows that TB does not always give rise to slow relaxation kinetics; it slows down the relaxation time of a positive feedback loop only when *ρ*_*u*_ ≪ *ρ*_*b*_, i.e. when the transcription rate in the repressed gene state is much smaller than that in the active gene state (Fig. 7(b)).

In the negative feedback case, we find no clear relationship between the relaxation rate and the two possible phases of the system (U or TB) (Fig. 7(c),(d)), which is in contradistinction to what we have observed in the positive feedback case. We emphasize here that while Fig. 7 displays the simulation results in the non-cooperative case, simulations in the cooperative case bear out the same conclusions.

Next we seek to understand the relationship between the dynamical phases of the system and the protein mean. In Fig. 8, we use heat maps to investigate how the size of the steady-state protein mean depends on the parameters *L, ρ*_*u*_, *ρ*_*b*_ and *B* for positive feedback loops (Fig. 8(a),(b)) and for negative feedback loops (Fig. 8(c),(d)). The regions of parameter space where each of the three phases (U, SB and TB) manifests are also shown. The most notable observation is that the regions of parameter space with the highest and lowest protein mean are also those where TB and U manifest, respectively. In addition, SB also manifests in the region where the protein mean is large.

**Fig. 8.**
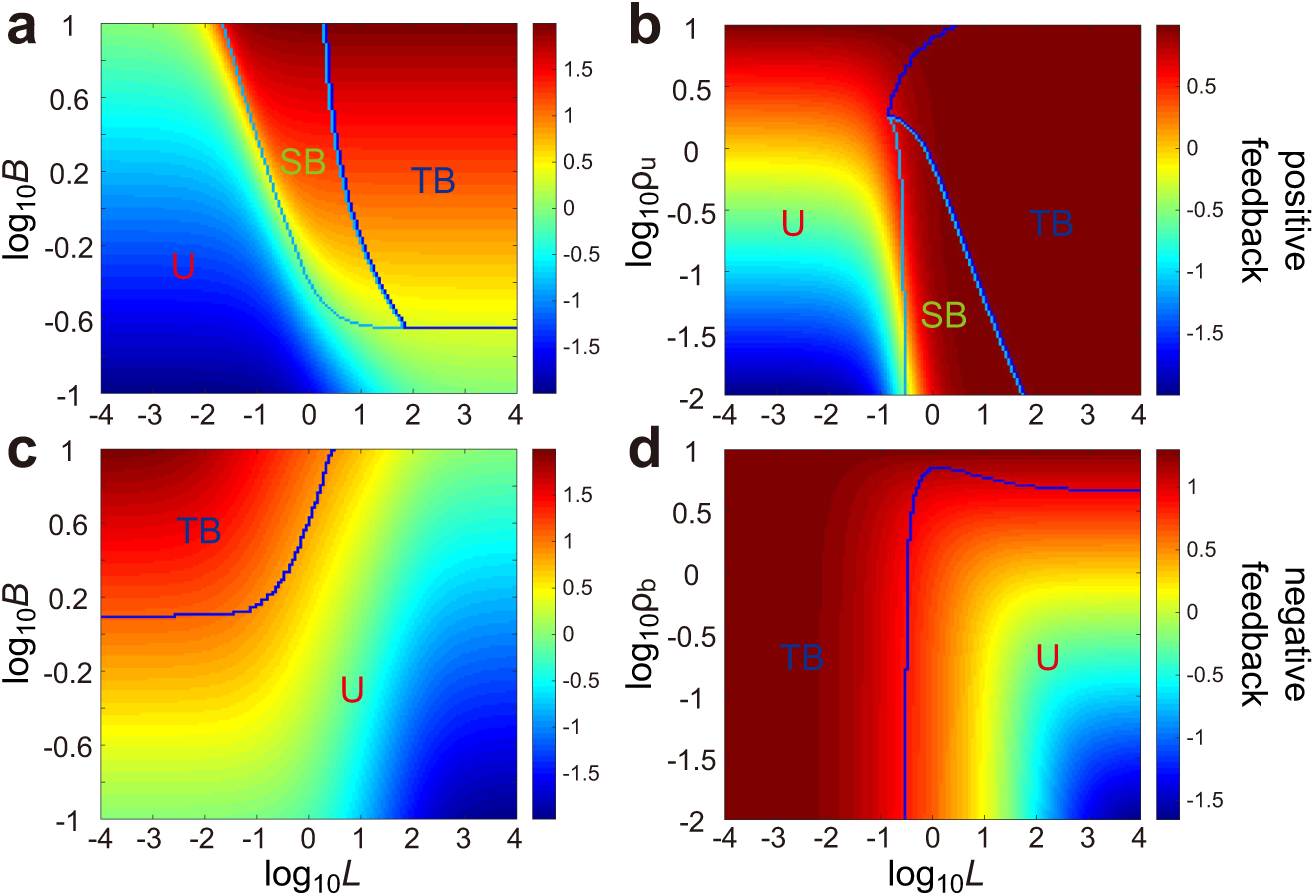
Steady-state protein mean for auto-regulated bursty gene expression. (a) Heat plot shows the steady-state protein mean log_10_⟨*n*⟩ as a function of the feedback strength *L* and the mean protein burst size *B* for positive feedback loops. (b) Heat plot shows the steady-state protein mean as a function of the strength *L* and the transcription rate *ρ*_*u*_ in the unbound gene state for positive feedback loops. (c) Heat plot shows the steady-state protein mean as a function of the feedback strength *L* and the mean protein burst size *B* for negative feedback loops. (d) Heat plot shows the steady-state protein mean as a function of the feedback strength *L* and the transcription rate *ρ*_*b*_ in the bound gene state for negative feedback loops. The model parameters in (a)-(d) are chosen to be the same as in the four subfigures of Fig. 7. Different phases (U, TB, and SB) are marked on each figure to show the relationship between the steady-state protein mean and the dynamical phases.

The connection between TB/SB and high protein mean regions can be understood as follows. Recall that a positive auto-regulatory gene circuit exhibits TB/SB when *L, B* ≫ 1. Now when *L* ≫ 1, the gene is mostly in the bound state which has a larger transcription rate than the unbound state. If both *L* and *B* are large, then the mean protein number at the steady state is also large. Similarly, recall that a negative auto-regulatory gene circuit exhibits TB when *L* ≪ 1 and *B* ≫ 1. When *L* ≪ 1, the gene is mostly in the unbound state which has a larger transcription rate than the bound state. If *L* is small and *B* is large, then the mean protein number at the steady state is also large. Therefore, in both positive and negative feedback cases, it is clear that the occurrence of TB/SB is closely related to the steady-state protein mean. While Fig. 8 shows simulation results in the non-cooperative case, simulations including cooperativity lead us to the same observations.

## 6 Summary and Discussion

In this paper, starting from a stochastic model of a bursty auto-regulating gene with cooperative protein-gene interactions, we used the multiscale averaging method to derive a reduced stochastic model describing protein dynamics in fast switching conditions. This model was then solved exactly in steady state and in time. The time-dependent solution is expressed in terms of the eigenvalues of the generator matrix of the reduced model which can be evaluated either numerically or else can be obtained from an approximate theory valid in the limit of fast or slow protein binding. The time-dependent solution was shown to excellently agree with numerical simulations using FSP and were used to identify six different dynamical phases of the system. The three main dynamical phases of the stochastic model are associated with the following types of time-evolution: (i) the protein distribution remains unimodal at all times (unimodality); (ii) the protein distribution becomes bimodal at intermediate times and then reverts back to being unimodal at long times (transient bimodality) and (iii) the protein distribution switches to being bimodal at long times (stationary bimodality). For each of these phases, the deterministic model can show either monostable or bistable behaviour at long times, hence implying the existence of six dynamical phases. If the deterministic and stochastic models share the same dynamic behaviour then noise is not important, whereas the opposite is true if their dynamic behaviours are contrasting. Out of the six phases, we find that only in two, noise is not important. Transient bimodality has no deterministic counterpart and hence noise plays a central role behind its manifestation.

We investigated the relationship between the dynamical phases and the transcription rates, the ratio of protein binding to unbinding rates, the relaxation rate and the mean protein number. While positive feedback loops display all six phases for sufficiently high cooperativity, negative feedback loops display only two. The most eclectic of these phases, namely the two phases which lead to transient bimodality (a purely noise-induced phenomenon), manifest provided that the translational burstiness in protein expression is large and protein binding to the gene is much faster (slower) than unbinding for positive (negative) feedback loops. Furthermore, for positive feedback loops, we used the theory to estimate the observation time window where transient bimodality occurs and showed that the phenomenon is associated with regions of parameter space where the mean protein numbers and relaxation times are large. In contrast, for negative feedback loops, we showed that there is no clear relationship between transient bimodality and the relaxation time but it is also associated with regions of parameter space where the mean protein numbers are high. We also demonstrated that in most biologically relevant cases, as the feedback strength increases, a positive feedback loop undergoes two stochastic bifurcations from the unimodality phase to the stationary bimodality phase and then to the transient bimodality phase, while a negative feedback loop undergoes only one stochastic bifurcation from the transient bimodality phase to the unimodality phase.

Our work thus advances our knowledge of the conditions under which it is possible to observe two seemingly different subpopulations (each associated with a mode of the protein distribution) in a population of identical cells. In particular, while it is currently thought that the transient appearance of two subpopulations can be due to a temporally varying stimulus in gene circuits with slow promoter switching (see Fig. 5 of Ref. [34]), we have shown that no such stimulus is needed in the presence of fast switching conditions (transient bimodality in phases 3 and 4). Furthermore we have shown that this mechanism does not rely on cooperativity and is enhanced when protein expression is sufficiently bursty, that is when many proteins are produced from a single mRNA copy, a fairly common scenario in eukaryotic cells because of the long lifetimes of eukaryotic mRNA and large translation rates [35, 36].

For simplicity and analytical tractability, we have here not included any explicit description of cell cycle effects. While we have an implicit effective description of protein dilution due to cell division, via the effective protein decay rate, it has recently been shown that in some parameter regimes, this type of model cannot capture the stochastic dynamics predicted by models with an explicit description of the cell cycle [37]. We hence anticipate that the inclusion of cell division and DNA replication may alter the time-evolution of protein distributions and may even introduce novel dynamical phases hitherto undescribed. These effects are currently under investigation.

## Acknowledgements

C. J. acknowledges support from the NSAF grant in National Natural Science Foundation of China (NSFC) with grant No. U1930402. R. G. acknowledges support from the Leverhulme Trust (RPG-2018-423). We thank James Holehouse and Kaan Ö cal for careful reading of the manuscript.

## Appendix

### A. Derivation of the deterministic rate equations

The CME of the auto-regulatory gene expression model is given by Eq. (2). Here we derive its mean-field approximation, i.e. we derive equations for the mean protein number and for the probability of the gene being in the bound or unbound state, under the assumption that the protein number is large and its fluctuations are negligible. Let 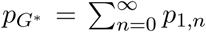 denote the probability of the gene being in the bound state and let 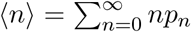 denote the mean of the protein number. If we interchange the order of the two sums, we obtain (when all terms are nonnegative, it follows from Fubini’s theorem that we can always interchange the order of the two sums)

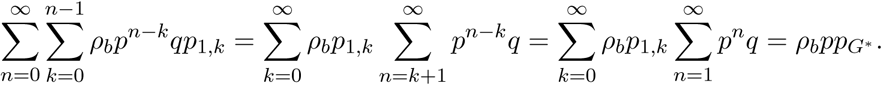

Moreover, it is easy to verify that

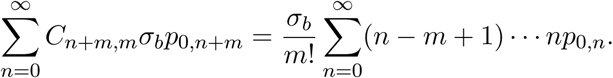

Combining the above two equations, it follows from of the CME that the evolution of *p*_*G\**_ is governed by the ordinary differential equation:

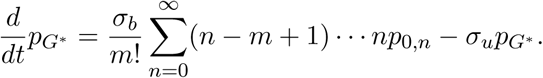

On the other hand, if we interchange the order of the two sums, we obtain

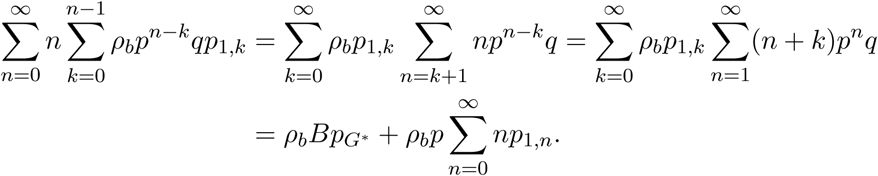

Moreover, it is easy to see that

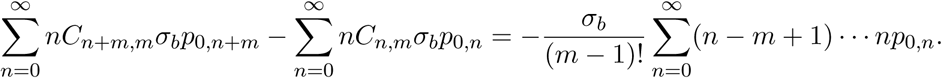

Combining the above two equations, it follows from of the CME that the evolution of ⟨*n*⟩ is governed by the ordinary differential equation:

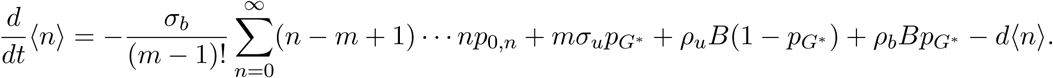

By the mean-field approximation, we assume that the protein number is large and its fluctuations are negligible, i.e. *n* ≈ *n* − 1 ≈ … ≈ *n* − *m* + 1 ≈ ⟨*n*⟩. This shows that

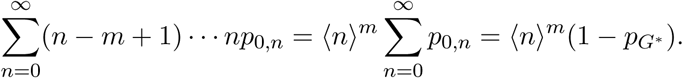

Thus, under the mean-field approximation, the evolution of *p*_*G\**_ and ⟨*n*⟩ is governed by the following coupled set of ordinary differential equations:

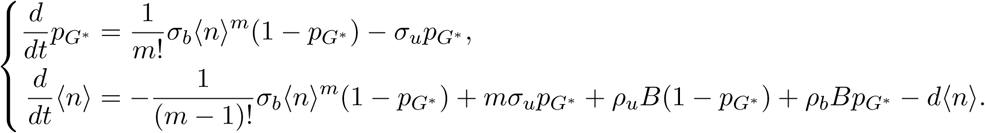

These are the deterministic rate equations given by Eq. (13) in the main text.

### B. Approximate eigenvalues for the case of slow protein binding

Our analytic solution for the time-dependent protein distribution, Eq. (18), depends on all eigenvalues of the generator matrix *Q*. In general, it is very difficult to compute these eigenvalues analytically. However, this can be done in two special cases: protein binding is much slower or much faster than protein unbinding.

We first focus on the case of *L* ≪ 1. In this case, protein binding is much slower than protein unbinding. Since the effective transcription rate *c*_*n*_ and effective protein decay rate *d*_*n*_ have the approximations given in Eq. (9), the generator matrix of the reduced model is then given by

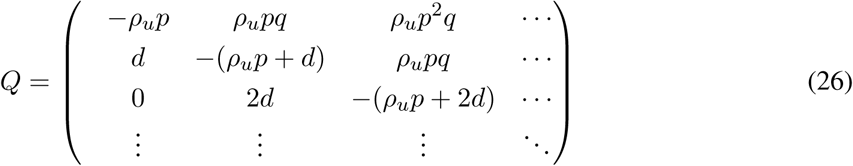

Recall that an eigenvalue-eigenvector pair (*λ, v*) of *Q* is related by the characteristic equation *vQ* = *λv*, which can be written in components as

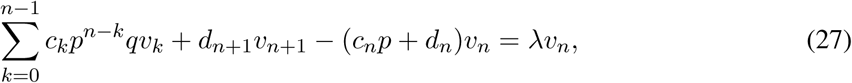

where we normalize the eigenvector *v* = (*v*_*n*_) so that *v*_0_ = 1. Using the approximation given in Eq. (9), the characteristic equation can be rewritten as

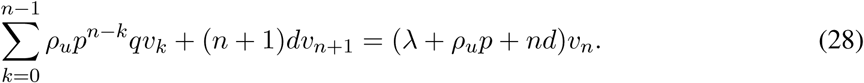

To proceed, we define the function *f* (*z*) to be the generating function of the eigenvector:

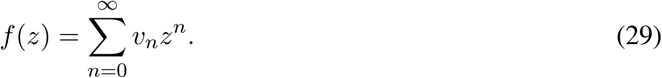

Since *v*_0_ = 1, we have *f* (0) = 1. Then Eq. (28) can be converted into the ordinary differential equation:

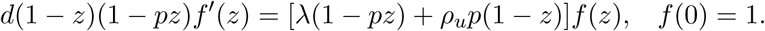

The solution of this equation is given by

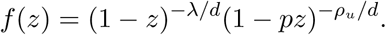

We next make a crucial observation that the components of the eigenvector *v* = (*v*_*n*_) must decay exponentially with respect to *n* when the system is ergodic [38]. In other words, *v*_*n*_ has the following approximation when *n* ≫ 1:

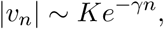

where *K* is a constant and *γ >* 0 describes the decay rate of *v*_*n*_ with respect to *n*. Thus, we have

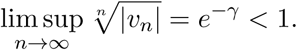

This shows that the convergence radius of the power series given in Eq. (29) must be greater than 1 and thus the generating function *f* (*z*) must be holomorphic on the unit circle. Here *f* (*z*) is the product of two terms, (1 − *z*)^−*λ/d*^ and 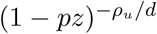. Since *p <* 1, the second term must be holomorphic on the unit circle. On the other hand, it is easy to see that the first term is holomorphic on the unit circle if and only if −*λ/d* is a nonnegative integer. This imposes a strong constraint on possible eigenvalues. Using this constraint, all the approximate eigenvalues of *Q* are given by

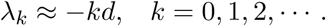

Substituting these approximate eigenvalues in Eq. (18) gives the approximate time-dependent protein distribution for slow protein binding conditions.

### C. Approximate eigenvalues for the case of fast protein binding

We consider the case of *L* ≫ 1, i.e. protein binding is much faster than protein unbinding. In this case, the effective transcription rate *c*_*n*_ and effective protein decay rate *d*_*n*_ have the approximation given in Eq. (10). Since *d*_*n*_ = (*n* − *m*)*d* for any *n* ≥ *m*, we have *d*_*m*_ = 0 and thus the generator matrix

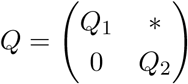

is an upper triangular block matrix, where the upper-left block is given by

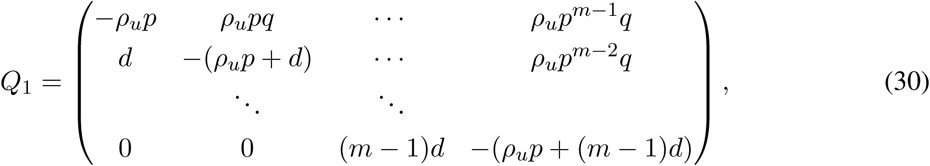

and the lower-right block is given by

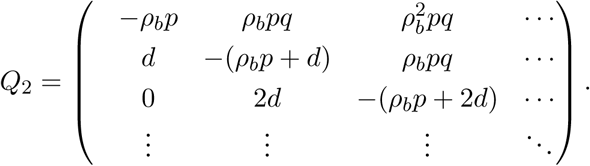

Therefore, the eigenvalues of *Q* are composed of the eigenvalues of both *Q*_1_ and *Q*_2_. Using elementary transformations, it is easy to prove that the eigenvalues of *Q*_1_ are given by the roots of the polynomial equation *u*_*m*_(*z*) = 0, where *u*_*m*_(*z*) is the polynomial defined in Eq. (16). Using the approximation given in Eq. (10), the polynomial *u*_*m*_(*z*) is defined recursively by

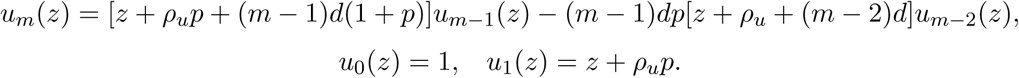

Since *u*_*m*_(*z*) is a polynomial of degree *m*, it has *m* zeros, say *x*_1_, …, *x*_*m*_. Moreover, we note that *Q*_2_ is exactly the matrix defined in Eq. (26) with *ρ*_*u*_ replaced by *ρ*_*b*_. Therefore, all the eigenvalues of *Q*_2_ are given by 0, −*d*, −2*d*, −3*d*, …. Then all the eigenvalues of *Q* are approximately given by

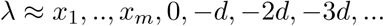

In particular, in the non-cooperative case of *m* = 1, we have *x*_1_ = −*ρ*_*u*_*p* and thus all the approximate eigenvalues of *Q* are given by

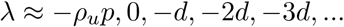

In the cooperative case of *m* = 2, we have

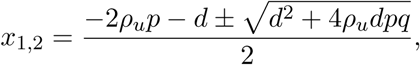

and thus all the approximate eigenvalues of *Q* are given by

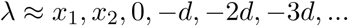

Substituting these approximate eigenvalues in Eq. (18) gives the approximate time-dependent protein distribution for fast protein binding conditions.

### D. Proof of a non-trivial equality

Here we shall give the proof of Eq. (22). In the main text, we have proved that

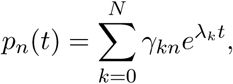

where

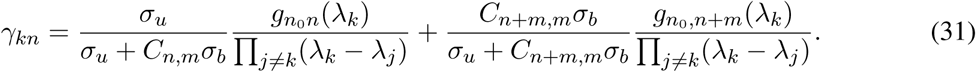

When *L* ≫ 1, we have computed all the approximate eigenvalues of *Q* in Appendix B. In particular, the first nonzero eigenvalue *λ*_1_ must satisfy

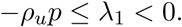

When *m* = 1, this is clear because the first nonzero eigenvalue is given by

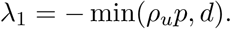

When *m* ≥ 2, it is easy to prove that the above inequality also holds. This clearly shows that

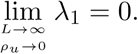

Since we have assumed that the initial protein number is zero, we have *n*_0_ = 0 and thus

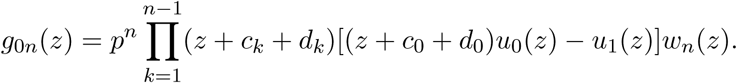

When *L* ≫ 1, we have *c*_0_ = *ρ*_*u*_ and *d*_0_ = 0. Since *u*_0_(*z*) = 1 and *u*_1_(*z*) = *z* + *c*_0_*p*, we have

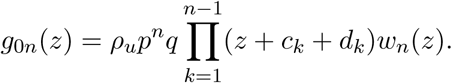

Since *λ*_0_ = 0 and *λ*_1_ ≪ 1, we have

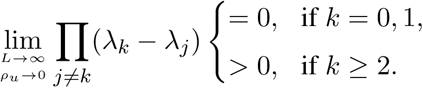

Combining the above two equations shows that for any *k* ≥ 2,

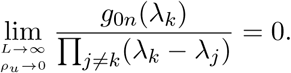

Inserting this equation into Eq. (31) gives Eq. (22).

